# Double stranded RNA sensing drives interferon silencing in early development

**DOI:** 10.1101/2025.03.10.642424

**Authors:** Jeroen Witteveldt, Zicong Liu, Ana Ariza-Cosano, Christian Ramirez, Jessica L. Walters, Pilar G. Marchante, Lars Maas, Alasdair Ivens, Toma Tebaldi, Sara R. Heras, Hendrik Marks, Sara Macias

## Abstract

In early mammalian development, the type I interferon (IFN) response is inactive, only becoming functional after gastrulation. As a result, the totipotent and pluripotent embryonic stages remain highly susceptible to pathogens, including viruses. Here, we show that pluripotent mouse embryonic stem cells (mESCs) suppress the RIG-I-like receptor sensing pathway by silencing the expression of the dsRNA sensor MDA5. We show that this silencing is required to avoid the recognition of dsRNAs from endogenous origin, which only accumulate in mESCs. Reintroducing MDA5 results in recognition of these endogenous dsRNAs, and activation of the IFN response through IRF3. The production of IFN alters the differentiation ability of mESCs, as well as the pluripotency gene expression program, as shown by epigenetic, transcriptomic and proteomic analyses. These findings are conserved in zebrafish, where MDA5 is also expressed at later stages of development. Similarly, zebrafish lack early-stage IFN activation and premature IFN signalling results in developmental defects. Altogether, we conclude that silencing the RIG-I-like receptor pathway during early development is the widely conserved and is required to prevent aberrant immune recognition of endogenous dsRNAs, safeguarding normal development.

## Introduction

Type I interferons (IFN) orchestrate the innate immune response against viruses. IFNs are produced by infected cells, typically upon sensing viral nucleic acids. Viral cytoplasmic DNA is sensed by the cyclic GMP-AMP synthase, cGAS and viral RNAs are sensed by the retinoic acid-inducible gene I (RIG-I)-like receptor (RLR) family of proteins (^1,2^). This family has three members, RIG-I, melanoma differentiation-associated protein 5 (MDA5) and laboratory of genetics and physiology (LGP2). RIG-I recognises uncapped short double stranded RNAs (dsRNAs), while MDA5 recognises long dsRNA molecules. Although LGP2 binds a variety of RNA moieties, the absence of CARD domains prevents downstream signalling. LGP2 regulates the activity of the other two members in the family; preventing RIG-I activation, while promoting MDA5 activity (^2^). Upon sensing of viral nucleic acids, several families of transcription factors are activated, including the interferon regulatory factors 3 and 7 (IRF3/7), which drive type-I IFN expression, and the nuclear factor-kappa B (NF-kB), which drives pro-inflammatory cytokines production, such as TNF-α (^3^). Despite high levels of homology, IRF3 is highly expressed in most cells, whereas IRF7 has very low basal expression levels. IRF7, which itself is an interferon stimulated gene (ISG), increases expression upon IFN activation, resulting in a positive feedback loop ensuring sustained levels of IFN expression (^4,5^). Besides type I interferons, IRF3 also induces expression of other immune-related genes, suggesting that it has a broader range of targets compared to IRF7 (^6–8^). Once produced and secreted, IFNs act in an autocrine and paracrine manner by binding IFN receptors (IFNAR1/2) and inducing the expression of hundreds of ISGs via JAK-STAT signalling (^9^). In addition to cytoplasmic sensors, the Toll-like receptors (TLRs), located on cell surfaces or endosomes can also recognise viral nucleic acids and activate the type I IFN response (^10^).

Healthy mammalian cells do not typically accumulate significant amounts of endogenous dsRNAs, as they have evolved a number of mechanisms to prevent their accumulation, (^11,12^). However, in certain disease settings, such as cancer or autoimmune disorders, mutations in important RNA processing enzymes, epigenetic regulators or nucleic acids sensing factors result in activation of the IFN response upon sensing endogenous dsRNAs from nuclear or mitochondrial origin, (^13,14^).

There is mounting evidence that transposable elements also represent a natural source of dsRNA, due to their repetitive nature and multiple insertions in sense or antisense orientation (^6–8^). However, most somatic cell types silence transposon expression to prevent the potential deleterious consequences of novel insertions. Interestingly, ESCs and the early stages of development are unique in terms of transposon activity. Contrary to somatic cells, the different stages of early embryonic development are characterised by waves of expression of specific TE families, including LINE-1 in the mouse 2-cell stage, or MERVL during early mouse development (^15^). In part, this is a consequence of the profound epigenetic reprogramming that embryonic cells undergo during the first divisions to acquire a totipotent or pluripotent identity, and that typically represses TE activity. As an adaptation, embryonic cells exploit TE, by co-opting TE promoters and TE-derived RNAs to execute essential functions during development _(_^16–19^).

ESCs and early development are also special in terms of immune defence. We and others have previously shown that the IFN response is inactive in pluripotent embryonic stem cells (ESCs) and early stages of mammalian development. Both mouse and human ESCs are incapable of producing IFNs when challenged with viruses or viral DNA/RNA mimics (^20–23^). Similarly, mouse embryonic teratocarcinoma cells, which retain certain pluripotency traits, and human induced pluripotent stem cells (iPSCs), are also incapable of IFN production (^23,24^). The ability to synthesise IFNs is acquired after differentiation, once cells have lost pluripotency (^20–23^). In agreement, *in vivo* data show that the ability to produce IFNs in mice is acquired after gastrulation (day 8 post-fertilisation) (^25^). These findings lead to the hypothesis that the IFN response and pluripotency are two incompatible cellular processes.

The ensuing question is why cells in early development, including ESCs, have sacrificed the type I IFN response if this renders them more susceptible to infections. Despite the existence of alternative antiviral pathways in ESCs, we and others have shown that for specific viruses, ESCs are more permissive than somatic cell types, arguing against a generalised intrinsic resistance of these cells to all viruses (^22,26–30^). Within the inner cell mass, ESCs may be physically protected from viruses by the trophectoderm, but also immunologically, as trophectoderm cells are immune competent and can stimulate the inner cell mass antiviral response by producing IFNs (^31–33^). Although attenuated, ESCs can respond to exogenous IFNs, suggesting that maternal or placental IFNs could have protective properties during development (^22,23,34,35^). For instance, constitutive expression of the type I IFN, IFN-ε, by the maternal reproductive system has been shown to provide protection against viral infection (^36^). To gain a better understanding of the mechanisms and rationale for the absence of the type I IFN response in early development, we employed pluripotent mESCs and zebrafish as models. We observed a silencing of the dsRNA sensing pathway in both models. mESCs supress the type I IFN response by silencing the expression of the dsRNA sensor MDA5. Unlike somatic cells, mESCs accumulate endogenous dsRNAs, which could induce the IFN response through MDA5 sensing. As a consequence, the introduction of MDA5 in mESCs leads to activation of the IFN response in the absence of infection. The inappropriate triggering of the IFN pathway is highly detrimental to mESCs, and impairs their differentiation capacity and pluripotency maintenance, as shown by epigenetic, transcriptomic and proteomic analyses. Silencing of the dsRNA sensing mechanisms is likely a conserved evolutionary strategy amongst jawed vertebrates. In zebrafish MDA5 expression is low during the early stages of development, and premature activation of the IFN response also results in significant developmental defects. Our current hypothesis is that the gene expression program necessary for early development is inherently incompatible with an active IFN response. TE expression is crucial for normal development but can result in the formation of dsRNAs that could trigger an IFN response through MDA5. Our results show that IFNs disrupt normal development, due to conflicts between IFN-associated transcription factors and the chromatin landscape in early development. Therefore, embryos must suppress the IFN response by downregulating dsRNA sensing, which, as a trade-off, increases their susceptibility to infections.

## Material and Methods

### Ethics statement

All experiments performed with zebrafish at the University of Granada comply with national and European Community regulations for the use of animals in experimentation and were approved by the ethical committees of the University of Granada and the Junta de Andalucía.

### Mouse and zebrafish datasets and analysis

Sequencing data from mouse embryonic stem cell differentiation studies were obtained from the Geo Expression Omnibus data repository (^37^, GEO125413) and ArrayExpress data collection (^38^, E-MTAB-4904). FastQ files were extracted from the repositories and quality assessed using FASTQC (v0.11.8). Cutadapt (v3.4) was used to trim the sequences using the command [cutadapt -a AGATCGGAAGAG -A AGATCGGAAGAG -j 0 -m 50 -o tr.$fq1 -p tr.$fq2 $fq1 $fq2 > tr.$sample.log]. Sequences were not collapsed within each sample, and duplicates were retained. Alignment of reads (--very-sensitive -p 24 --no-mixed --no-discordant --no-unal -x $db -1 tr.$fq1 -2 tr.$fq2 2>> $bt2log) to the REL104 reference set was performed using hisat2 (v 2.2.1). Alignment rates were typically 90% of input reads per sample. Alignments were stored in sorted, indexed BAM files. Raw tag counts per gene in each sample were extracted from the BAM files using the gtf file in combination with featureCounts function of Rsubread (featureCounts(files = bamfiles, annot.ext = ’Mus_musculus.104.gtf.gz’, isGTFAnnotationFile = TRUE, countMultiMappingReads = TRUE, allowMultiOverlap = TRUE, isPairedEnd = TRUE, nthreads = 24, minOverlap = 10)), and were normalised as part of the DESeq2-based groupwise comparisons: no conversions (e.g. reads per kb of gene, reads per million, etc.) were performed. The zebrafish dataset was obtained from the Geo Expression Omnibus data repository (^39^, GSE106430). Raw reads were aligned to the complete assembly of zebrafish genome GRCz11 and quantified according to NCBI RefSeq *Danio rerio* Annotation Release 106 (June 2, 2017) using STAR v2.7.11b (^40^) and quantified according to NCBI RefSeq Danio rerio Annotation Release 106 (June 2, 2017) using featureCounts function of Subread v2.0.8.

### Cells and differentiation protocols

The mouse embryonic stem cell line v6.5 was obtained from ThermoFisher (MES1402) and cultured in Dulbecco’s modified Eagle Medium (DMEM, ThermoFisher) supplemented with 15% heat-inactivated foetal calf serum (ThermoFisher), 1X Minimal essential amino acids (ThermoFisher), 2 mM L-glutamine, 10^3^ U/ml of LIF (Stemcell Technologies) and 50 µM 2-mercaptoethanol (ThermoFisher). Stem cells were grown on plates coated with 0.1% gelatine, detached using 0.05% Trypsin (ThermoFisher). The mouse microglial BV-2 cells were cultured in Dulbecco’s modified Eagle Medium (DMEM, ThermoFisher) supplemented with 10% heat-inactivated foetal calf serum (ThermoFisher) and 2 mM L-glutamine. Cells were incubated at 5% CO_2_ at 37°C.

Mouse embryonic stem cells were differentiated using hanging droplets and retinoic acid to induce embryoid body formation and differentiation as described before (^41^). Differentiated cells were collected and RNA was extracted using Tri-reagent (Sigma-Aldrich) and processed for real time PCR as described below.

For poly(I:C) treatment, differentiated cells were collected and seeded in 24 well plates, transfected with 2.5 µg poly(I:C) using lipofectamine 2000 and incubated for 6 hours before collecting the cells in Tri-reagent. For exogenous IFN-β treatment, differentiated cells were collected and seeded in 24 well plates and incubated with 10 or 100 pg/ml of mouse IFN-β (Bio-techne, 8234-MB-010) for 4 hours before collecting the cells in Tri-reagent. To assess the impact of the IFN response during differentiation a modified differentiation protocol was used. V6.5 and MDA5-overexpressing cells were pre-treated with doxycycline for 8 hours before a single-cell suspension of 5 × 10^5^ cells/ml was prepared in medium without LIF and in the presence or absence of doxycycline. Twenty µl drops were pipetted on the inside of the lid of a 10 cm petri dish and incubated upside-down at 37°C, 5% CO2 for 24 hr. The embryoid bodies were consequently washed from the lids, pelleted by centrifugation and RNA extracted using Tri-reagent.

### RNA extraction and real time PCR

Total RNA was extracted using Tri-reagent (MilliporeSigma), and cDNA was synthesized using M-MLV (Promega) or the High-Capacity cDNA Reverse Transcription Kit (Applied Biosystems) in accordance with the manufacturer’s instructions. qPCR reactions were performed using GoTaq qPCR mastermix (Promega) or qPCRBIO sygreen (PCR Biosystems) using previously published primers (Supplementary file 5) on a QuantStudio 5 (ThermoFisher) or Azure Cielo 6 (Azure biosystems). Data was analysed using Quantstudio Design & Analysis software or Azure Cielo Manager software. Differences were analysed by single factor ANOVA to calculate significant differences amongst comparisons, followed by an F-test for variance and appropriate two tailed t-test (*) p-val≤0.05, (**) p-val≤0.01, (***) p-val≤0.001.

### Western blot assay

Whole-cell extracts were collected in RIPA buffer containing protease inhibitors followed by sonication and centrifugation for clarification of extracts. Extracts were quantified using the BCA assay, mixed with LDS sample buffer (Invitrogen) and Sample reducing agent (Invitrogen) and run on 4-12% pre-cast gels (ThermoFisher). Proteins were transferred to nitrocellulose membrane using semi-dry transfer (iBlot2, ThermoFisher). Membranes were blocked for 1 hr at room temperature in PBS-T (0.1% Tween-20) and 5% milk powder before overnight incubation at 4°C with primary antibody. Antibodies used were anti-mouse HRP (Bio-Rad), anti-FLAG (M2, Merck), anti-tubulin (CP06, Calbiochem). Proteins bands were visualised using ECL (Pierce) on a Bio-Rad ChemiDoc imaging system. Protein bands were quantified using ImageJ (v1.53q) software and expression levels calculated normalized to tubulin.

### J2 flow cytometry

Cells were dissociated using 0.05% Trypsin, washed in PBS and resuspended in FACS buffer (PBS with 1% FBS). Cells were pelleted and resuspended in Fixation buffer (420801, BioLegend) and incubated for 15’ at 4°C, washed twice with Intracellular Staining Permeabilization Wash Buffer (421002, BioLegend) and stained overnight with the anti-dsRNA antibody J2 (English & Scientific Consulting) in Intracellular Staining Permeabilization Wash Buffer. After washing, cells were incubated with anti-mouse Alexa Fluor 647 (ThermoFisher) for one hour at room temperature and washed three times with Intracellular Staining Permeabilization Wash Buffer before finally resuspending the cells in FACS buffer. Cells were analysed using a MACS Quant analyzer 10 (Miltenyi), and data were processed using FlowJo software (Treestar).

### Stable cell lines

Plasmids containing the sequences of mouse MDA5 (GE-healthcare, MMM1013-202762875) and 3xFLAG (pcDNA3.1-3xFLAG) were used to amplify Gibson assembly fragments to construct the MDA5 sequence with a 3xFLAG tag at the N-terminal end. A plasmid containing the MAVS sequence (GE-healthcare, MMM1013-202764911) was used as template to directly amplify the MAVS open reading frame with specific restriction sites and a N-terminal FLAG tag (for oligonucleotides, see Supplementary file 5). The amplified and digested fragments were ligated into the pLenti-GIII-EF1a plasmid for constitutive expression and/or purification, and the pCW57-MCS1-P2A-MCS2 (89180, Addgene) plasmid, in which we replaced the hPGK promotor with the EF1a promotor for optimal expression in mESCs. For the ΔCTD mutant of MDA5, a reverse primer containing a stop codon and specific restriction site was used to amplify a fragment that was ligated into the same pLenti-GIII-EF1a and pCW57-MCS1-P2A-MCS2 plasmids. Verified plasmids containing the genes of interest were transfected in mESCs using Lipofectamine 2000 and selected with the appropriate antibiotic. Clonal cell lines were isolated, expanded and tested for expression by qRT-PCR and Western blot. ShRNA sequences targeting IRF3, IRF7 and MAVS were designed using the Broad institute design tool (https://portals.broadinstitute.org/gpp/public/gene/search) and ligated into the pLKO.1 plasmid (24150, Addgene). After sequencing to verify to integrity, the plasmids were transfected in mESCs using Lipofectamine 2000 and selected with the appropriate antibiotic. Clonal lines were isolated, expanded and tested for knock down of the target gene by RT-qPCR (Supplementary file 5).

### MDA5 purification and poly(I:C) pulldown

HEK293T cells were transfected with the pLenti-GIII-EF1a plasmids expressing FLAG-MDA5 and FLAG-ΔCTD using Lipofectamine 2000, following manufacturer’s instructions. Cells were collected after 48h and resuspended in lysis buffer (20 mM HEPES-KOH pH 7.9, 100 mM KCl, 0.2 mM EDTA, 0.5 mM DTT, 02. mM PMSF, 5% glycerol, supplemented with Complete Protease inhibitors), followed by sonication. Lysates were incubated with pre-washed FLAG Dynabeads (Sigma-Aldrich, M8823) overnight. Beads were washed five times with lysis buffer, supplemented with 1M KCl, with an additional final wash with TBS (50 mM Tris pH 7.4, 150 mM NaCl). Purified MDA5 proteins were eluted using FLAG peptide (Sigma-Aldrich, F4799) (100 ug/ml final concentration) in TBS buffer, following manufacturer’s instructions. Eluted proteins were quantified by Coomassie staining and nanodrop. For poly(I:C) pulldowns, poly(C)-agarose beads (Sigma-Aldrich, P9827) were washed twice with 50 mM Tris pH 7.0, 200 mM NaCl, and resuspended in 50 mM Tris-pH 7.0, 50 mM NaCl. Poly(I) (Sigma-Aldrich, P4154) was dissolved in 50 mM Tris-pH 7.0, 150 mM NaCl to a concentration of 2 mg/ml. One part of poly(C) beads and two parts of poly(I) solution were mixed and rocked for 1h at 4C to form double-stranded RNA (dsRNA). Next, beads were pelleted, washed and resuspended in 50mM Tris-pH 7.0, 150 mM NaCl. Poly(C) beads only were used as a control (single-stranded RNA, ssRNA). For pulldowns, equilibrated dsRNA-beads were resuspended in binding buffer (50 mM Tris-pH 7.5, 150 mM NaCl, 1 mM EDTA, 1% NP-40). Equal volume of beads slurry was combined with an equal volume of MDA5 protein (300 ug of WT or ΔCTD), plus protease, phosphatase and 25 U RNAsin/ml. Reactions were incubated at 4C with gentle rotation for 1h, washed three times with binding buffer and resuspended in SDS-PAGE loading buffer for downstream analyses by western blot.

### Total RNA high-throughput sequencing and analysis

Total RNA was extracted from cells using Tri-reagent and the quality assessed on the Agilent 2100 Electrophoresis Bioanalyser Instrument (Agilent, G2939AA). RNA was quantified using the Qubit 2.0 Fluorometer (ThermoFisher, Q32866). DNA contamination was quantified using the Qubit dsDNA HS assay kit and confirmed to be < 6% (ThermoFisher, Q32854). Libraries for total RNA sequencing were prepared using the NEBNext Ultra 2 Directional RNA library prep kit (Illumina, E7760) and the NEBNext rRNA Depletion kit (Human/Mouse/Rat) (Illumina, E6310) according to the provided protocol. Sequencing was performed on the NextSeq 2000 platform (Illumina, 20038897) using NextSeq 2000 P3 Reagents (200 Cycles, 2×100bp) (20040559). The raw sequences from the two sequencing runs were combined prior to being quality assessed using FASTQC (v0.11.8), and html format outputs generated. Cutadapt (v3.4) was used to trim the sequences with the following command: cutadapt -a AGATCGGAAGAG -A AGATCGGAAGAG -j 0 -m 50 -o tr.$fq1 -p tr.$fq2 $fq1 $fq2 > tr.$sample.log. Sequences were not collapsed within each sample, and duplicates were retained. The murine genome (release 104, “REL104”) and gtf formatted annotation was obtained by ftp from ensembl (ftp.ensembl.org/pub/release-104/fasta/mus_musculus). Alignment of reads (--very-sensitive -p 24 --no-mixed --no-discordant --no-unal -x $db -1 tr.$fq1 -2 tr.$fq2 2>> $bt2log) to the REL104 reference set was performed using hisat2 (v 2.2.1. Alignments were stored in sorted, indexed BAM files. Raw tag counts per gene in each sample were extracted from the BAM files using the gtf file in combination with featureCounts function of Rsubread (featureCounts(files = bamfiles, annot.ext = ‘Mus_musculus.104.gtf.gz’, isGTFAnnotationFile = TRUE, countMultiMappingReads = TRUE, allowMultiOverlap = TRUE, isPairedEnd = TRUE, nthreads = 24, minOverlap = 10)), and were normalised as part of the DESeq2-based groupwise comparisons: no conversions were performed. Pairwise comparisons of sample groups were performed using the DESeq2 Bioconductor package, with alpha set at 1; as each sample group had 3 replicates, all groups were used. logFC values were shrunk using the ashr model in lfsShrink() and a significance threshold of p<0.01 (adjusted) was applied.

Gene ontology (biological process) analysis was performed using the ClueGo plugin in the Cytoscape package (^42^). Cell type predictions were performed using the Mouse Gene Atlas within the Enrichr package (https://maayanlab.cloud/Enrichr/) (^43^). Transcription factor predictions were done using the LOF dataset within the ShinyGO (0.81) package (^44^).

### Chromatin enrichment for proteomics (ChEP)

ChEP was conducted based on previously established protocols with slight modifications (^45,46^). Briefly, 1 × 10⁷ cells were crosslinked using 1% formaldehyde at room temperature (RT) for 10 minutes, followed by quenching with 0.25 M glycine. The cells were lysed in 1 ml ice-cold cell lysis buffer (25 mM Tris, pH 7.4, 0.1% Triton X-100, 85 mM KCl, and 1× Roche protease inhibitor) and centrifuged at 2,300 × g for 5 minutes at 4 ℃ to isolate the nuclei. After discarding the cytoplasmic fraction, the pellet was resuspended in 500 μl SDS buffer (50 mM Tris, pH 7.4, 10 mM EDTA, 4% SDS, and 1× Roche protease inhibitor) and incubated at RT for 10 minutes. The solution was diluted to 2 ml using urea buffer (10 mM Tris, pH 7.4, 1 mM EDTA, and 8 M urea) and centrifuged at 16,100 × g for 30 minutes at RT, repeating the step once. Subsequently, the pellet was resuspended with 2 ml SDS buffer and centrifuged at 16,100 × g for 30 minutes at RT. The resulting protein pellet was dissolved in 250 μl storage buffer (10 mM Tris, pH 7.4, 1 mM EDTA, 25 mM NaCl, 10% glycerol, and 1× Roche protease inhibitor) and sonicated using a Bioruptor (Diagenode) for five cycles (30 seconds on/off) to enhance solubility. Protein concentration was quantified via a Qubit protein assay (Invitrogen). For mass spectrometry (MS) analysis, 100 μg of protein extract was de-crosslinked at 95 ℃ for 30 minutes. Each timepoint included two biological replicates.

For proteomic analyses, de-crosslinked chromatin protein extracts (100 μg) were precipitated by adding 10 volumes of ice-cold 100% acetone and incubated at -20 ℃ for 15 minutes. The samples were centrifuged at 16,000 × g for 10 minutes in a pre-cooled centrifuge. The resulting protein pellets were washed with 500 μl ice-cold acetone, centrifuged, and the acetone was discarded. The pellets were resuspended in 100 μl urea buffer (8 M urea, 100 mM Tris-HCl, pH 8.0, 10 mM DTT) and incubated at room temperature for 20 minutes. Subsequently, 10 μl iodoacetamide was added, and the mixture was incubated in the dark for 15 minutes at room temperature. For protein purification and on-bead digestion (^47^), 5 μl of pre-washed carboxyl magnetic beads (Thermo Scientific) and 154 μl binding buffer (100% acetonitrile, ACN) were added, followed by incubation for 18 minutes at room temperature. The beads were washed twice with 180 μl of 70% ethanol and once with 200 μl of 100% ACN. Proteins were eluted and digested on beads in 100 μl of ammonium bicarbonate buffer (ABC) containing 3 μl trypsin (Promega) at 37 ℃ for 2 hours. After removing the beads, the solution was incubated at 37 ℃ overnight to complete digestion. The digested peptides were fractionated using strong anion exchange (SAX) into three fractions (^48^): flow-through (FT), pH 8 elution, and pH 2 elution. The pH 8 and pH 2 fractions were combined. Finally, peptides were enriched and desalted using Stage-Tip (^49^) desalting before liquid chromatography-tandem mass spectrometry (LC-MS/MS) analysis.

Proteomic data analysis was carried out following previously described protocols with minor modifications (^46^). Raw mass spectrometry data were processed using MaxQuant software (^50^) (v2.4.2) with searches performed against the mouse UniProt database (downloaded June 2017). Default parameters were used, with LFQ, iBAQ, and the “match between runs” feature enabled. Protein identification was validated against a decoy database generated within MaxQuant. Biological replicates were grouped to identify differentially expressed proteins. Data were filtered for three valid values in at least one group. Missing values were imputed using default settings in Perseus (^51^) (v2.0.11) based on the assumption that they were not detected because they were under or close to the detection limit. Differentially enriched proteins were determined using a two-tailed Student’s t-test (P < 0.05) with a fold change threshold of >2. Differential proteins at 8h, 24h, and 48h were excluded from the differential proteins at 0h. Downstream analyses, including data visualization, were performed using R software (version 4.4.1).

### Assay for Transposase-Accessible Chromatin using sequencing (ATAC-seq) and analysis

The ATAC (mitochondrial DNA-remove) protocol was adapted from previously established methods with modifications (^52,53^). A total of 10,000 cells were collected, washed with 1 mL of ice-cold PBS containing EDTA-free protease inhibitor, and centrifuged at 500 × g for 5 minutes at 4 ℃. The supernatant was completely removed, and the cell pellet was resuspended in 20 μL of transposase reaction mixture, consisting of 1×Tagment DNA buffer, 1 μL Tn5 enzyme, and 0.02% digitonin. The transposition reaction was carried out at 37 ℃ for 30 minutes with continuous agitation. Following transposition, DNA was purified and the ATAC-seq library was constructed using the KAPA HyperPrep Kit (KAPA Biosystems, 07962363001) and barcoded with NEXTflex DNA barcodes (Integrated DNA Technologies). Paired-end sequencing of the library was performed on an Illumina NextSeq 500 platform.

Sequence reads were processed using the seq2science pipeline (v1.2.2)(^54^). Paired-end reads were trimmed with fastp v0.23.2 with default options. Genome assembly GRCm39 was downloaded with genomepy 0.16.1. Reads were aligned with bwa-mem2 v2.2.1 with options ‘-M’. Afterwards, duplicate reads were marked with Picard MarkDuplicates v3.0.0. Before peak calling, paired-end info from reads was removed with seq2science so that both mates in a pair get used. Bam files were randomly sampled down to the reads number as the same as the lowest reads sample. Peaks were called with macs2 v2.2.7 with options ‘ --shift 75 --extsize 200 – nolambda’. Differential peaks were calculated by Diffbind (v3.12.0). Peak tracks and heatmaps were generated by Deeptools (v3.5.5). Peaks were annotated by ChIPseeker v1.38.0. GO analysis was conducted by clusterProfiler v4.10.1.

### Double-stranded RNA immunoprecipitations and analyses

Double stranded RNA was isolated using the dsRNA specific antibody J2 (English & Scientific Consulting). For each replicate, the cells from one confluent 10cm plate were collected, washed twice with cold PBS and lysed in 1 ml IP-buffer (50mM Tris pH 7.5, 150mM NaCl, 1mM EDTA, 1% Triton X-100) supplemented with protease inhibitors (S8820, Sigma-Aldrich). For each 1 ml of lysate, 10μl DNAse (M6101, Promega) and 2μl RNAse inhibitor (N2111, Promega) was added and incubated on ice for 30 minutes after which the lysate was spun at 13krpm, 4°C for 15’ and the supernatant collected.

To couple J2 antibodies to magnetic protein G beads, 50 μl of protein G slurry (88848, Thermo Scientific) was washed three times with cold IP buffer, resuspend in 100 μl IP buffer with 5 μg of J2 antibody and mixed at 4°C while rotating for 3 hours. After incubation, the beads were washed three times with IP buffer, mixed with the lysate and incubated at 4°C for 4 hours while rotating. The beads were washed three times 5 minutes at 4°C while rotating and tubes were changed once to prevent RNA carry over. After the last wash, the beads were resuspended in 250μl of 0.3M NaAc (pH 5.2) and RNA was extracted using Tri reagent LS according to the manufacturer’s protocol.

Total RNA was processed according to the protocol described in the ‘Total RNA high-throughput sequencing and analysis’ section. Sequencing reads were processed, aligned and counted according to the protocol described in the ‘Total RNA high-throughput sequencing and analysis’ section. Normalization (TMM method) and differential expression analysis between J2 IP and RNAseq gene counts were performed using the glmQLFit function from edgeR v. 4.2.2 R package, as described in ^55,56^. Transcripts were classified as enriched dsRNAs when J2 IP vs RNAseq log2FC > 0.75 and false discovery rate (FDR) < 0.05. Minimum free energy values were calculated via RNAfold from the ViennaRNA package v. 2.7.0 (^57^). Transcript sequences were obtained from GENCODE chromosome sequences (GRCm39) and Ensemble genome annotation (release 104, “REL104”). Transposable elements (TE) annotation was obtained from the Hammel Lab (https://labshare.cshl.edu/shares/mhammelllab/www-data/TEtranscripts/TE_GTF/) and overlap with gene coordinates was performed with findOverlaps function from the GenomicRanges v. 1.56.2 R package.

### Zebrafish experiments

Zebrafish wild-type strains AB/Tübingen/TAB (AB/Tu/TAB) were gifts from the zebrafish facilities at LARCEL (Málaga, Spain) and MPI-CBG (Dresden, Germany). The fish were maintained and bred under standard conditions (^58^). Wild-type zebrafish embryos were obtained by natural mating of AB/Tu and TAB zebrafish of mixed ages (5-18 months). Pairs were randomly selected from 20 males and 20 females and zebrafish embryos were staged as described before (^59^). After fertilization, eggs were collected and one-cell stage embryos were microinjected with 2 nL of 1 μg/μL poly(I:C) in PBS (^60^). Control embryos were injected with the same volume of PBS.

For whole-mount *in situ* hybridization, a partial region of Trim25 was cloned into pGEM-T using specific primers (Supplementary file 5), linearized with NcoI and *in vitro* transcribed using SP6 (P1085, Promega) to generate an antisense trim25 probe, which was labelled with digoxigenin. Three, six and 24 hours post fertilization and poly(I:C) injection, zebrafish embryos were fixed, hybridized, and stained using NBT/BCIP (^61^), imaged and analysed using ImageJ.

Total RNA was isolated from 50-60 embryos 24 hours post fertilization and poly(I:C) injection using Trizol (Invitrogen) according to the manufacturer’s protocol. One microgram of RNA was DNase I treated and purified via phenol/chloroform extraction. cDNA was synthesised using the High-Capacity cDNA Reverse Transcription Kit (Applied Biosystems) and used for qPCR (GoTaq qPCR Mix, Promega).

Poly(I:C) and PBS injected embryos of the same age were used to compare the differences in developmental defects due to their treatments. Embryos were classified based on severity of the development defects, where embryos were assigned class 1 defects if embryos exhibited failure to develop the head and class 2 defects if there were defects in trunk development.

## Results

### The expression of the dsRNA sensor MDA5 is developmentally regulated

ESCs fail to make IFNs when challenged with dsRNA, but they acquire this ability after differentiation (^20–22,62–65^). To identify which components of the IFN response may be responsible for the differential ability in producing IFNs, we compared the expression of genes involved in TLR signalling, RLR signalling and JAK/STAT signalling during stem cell differentiation. We used two different datasets, one of mouse ESCs differentiated to the three major embryonic lineages, ectoderm, mesoderm and endoderm (^38^), and a second, where mouse ESCs were differentiated to neuronal stem cells and neurons (^37^) (Fig. 1a and Extended data Fig. 1a,b). Although varying between lineages, we observed a consistent increase in the levels of expression of *Ifih1*, the gene encoding for the dsRNA sensor MDA5, and *Irf*7, which encodes for the transcription factor IRF7. The expression of both genes was negligible in ESCs and increased after differentiation (Fig. 1a and Extended data Fig. 1a,b). Both genes are involved in activating IFN production upon dsRNA stimulation, suggesting that the RLR pathway is silenced at multiple levels in ESCs. We confirmed these findings using *in vitro* differentiation assays of ESCs through induction of embryoid body formation followed by retinoic acid treatment (^41^). After 20 days of differentiation, the expression of pluripotency factors *Nanog* and *Pou5f1* was lost, while the expression of differentiation markers *Sox17* and *Foxa2* increased as expected. We observed that, upon differentiation, *Ifih1* was the most differentially expressed gene involved in the RLR signalling pathway, with a significant increase after differentiation (Fig. 1b). We confirmed that upon differentiation, and increased *Ifih1* expression, mESCs acquire the ability to respond to the dsRNA mimic, poly(I:C), as measured by RT-qPCR of IFNs (*Ifnb1*), *Tnfa* as well as of the ISGs, *Cxcl10* and *Ifih1* (Fig. 1c). As sensing type I IFN is integral to the overall IFN response, we also compared the response of ESCs and differentiated cells to exogenous IFN-β. ESCs displayed an attenuated response to exogenous IFN-β, which also increases upon differentiation, as shown by RT-qPCR analyses of a panel of ISGs (Fig. 1d). All these together confirmed that ESCs acquire the IFN response during differentiation, and this coincides with a significant upregulation of the dsRNA sensor MDA5 (*Ifih1*).

**Fig. 1.**
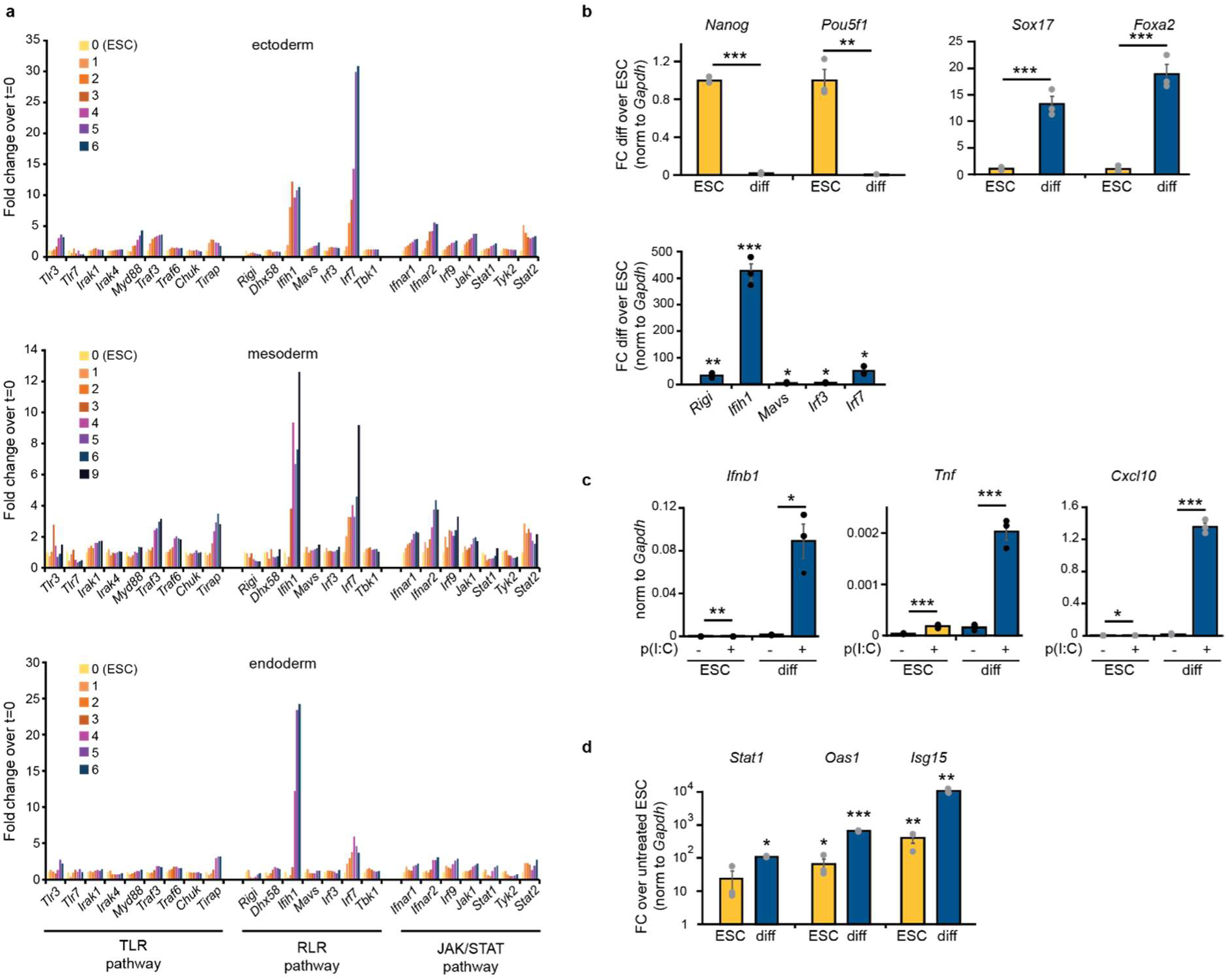
*Ifih1* expression is induced upon differentiation. (**a**) Expression of genes involved in TLR signalling, RLR signalling pathway and JAK/STAT signalling in mouse embryonic stem cells (mESCs) differentiating to ectoderm (top), mesoderm (middle) and endoderm (bottom), as obtained from RNA-seq. Samples were taken at successive days after starting differentiation (day 0 corresponds to ESCs, until day 6 or 9 of differentiation). Expression is calculated as normalised counts and relative to ESCs (day 0). (**b**) RT-qPCR analyses of embryoid body differentiated (Diff) and pluripotent mESCs. Data represent the average of three biological replicates ± the standard error of the mean (SEM). Single factor ANOVA was used to calculate significant differences amongst comparisons, followed by an F-test for variance and appropriate two-tailed t-test. (**C**) RT-qPCR analyses of *Ifnb1* and ISGs after dsRNA stimulation with poly(I:C) (p(I:C)) of ESCs and *in vitro* differentiated ESCs (as in **b**). Data represent the average of three biological replicates ± SEM. Single factor ANOVA was used to calculate significant differences amongst comparisons, followed by an F-test for variance and appropriate two-tailed t-test (**d**) ESCs and *in vitro* differentiated cells (as in **b**) were stimulated with exogenous IFN-β (100pg/ml), ISGs induction was quantified by RT-qPCR. Data represent the average of three biological replicates ± SEM. Single factor ANOVA was used to calculate significant differences amongst comparisons, followed by an F-test for variance and appropriate two-tailed t-test. (*) p-val≤0.05, (**) p-val≤0.01, (***) p-val≤0.001.

### Introduction of MDA5 in ESCs leads to developmental disruption

MDA5 is expressed at very low levels in ESCs, therefore we decided to explore the consequences of expressing MDA5 in these cells by generating ESC lines constitutively expressing MDA5. Although MDA5 expression was driven by a promoter known to be stable in mESCs (EF1a), after several passages, cells silenced the exogenous copy of MDA5 in an irreversible manner. To address this issue, we generated doxycycline-inducible ESCs lines expressing a FLAG-tagged form of MDA5. Two clones were selected for further characterisation, and MDA5 expression after doxycycline treatment was confirmed both by western-blot and RT-qPCR (Fig. 2a,b, respectively). Also in this set-up, we observed silencing of MDA5 after 24h, suggesting an adverse effect of long-term expression of MDA5. To further investigate the consequences MDA5 expression in ESCs, we performed a time course analysis by RNA-high-throughput sequencing (RNA-seq) after MDA5 expression (0, 4, 8, 24 hours after doxycycline induction) with two independent clonal cell lines compared to wild type ESCs (WT). Surprisingly, hundreds of genes were commonly differentially expressed in both MDA5-expressing cell lines compared to WT, with significant changes as early as 4 hours after induction (abs log2FC > 0.4; pval ≤ 0.05) (Supplementary Excel file 1). Gene ontology (GO) analysis of the shared differentially expressed genes showed that both upregulated and downregulated genes were associated with development, morphogenesis, and differentiation, confirming the profound rewiring of the developmental programme when MDA5 is expressed (Fig.2c and Extended data Fig. 2a,b, Supplementary Excel file 2). More specifically, the upregulated genes were associated with terms involving nervous system development. This was confirmed using the Mouse Gene Atlas to predict cell types based on differential gene expression data. Upregulated genes were associated with cell types from the nervous system, suggesting that MDA5 induction in ESCs results in a ‘neuronal-like’ gene expression profile (Fig. 2c,d). Conversely, downregulated genes were associated with genes typically expressed in ESCs and embryonic fibroblasts, suggesting that MDA5 induces the loss of the embryonic stem cell features (Fig. 2c,d). We conclude that the expression of MDA5 causes a profound disruption of the gene expression programme of ESCs, leading to an expression profile associated with differentiating cells.

**Fig. 2.**
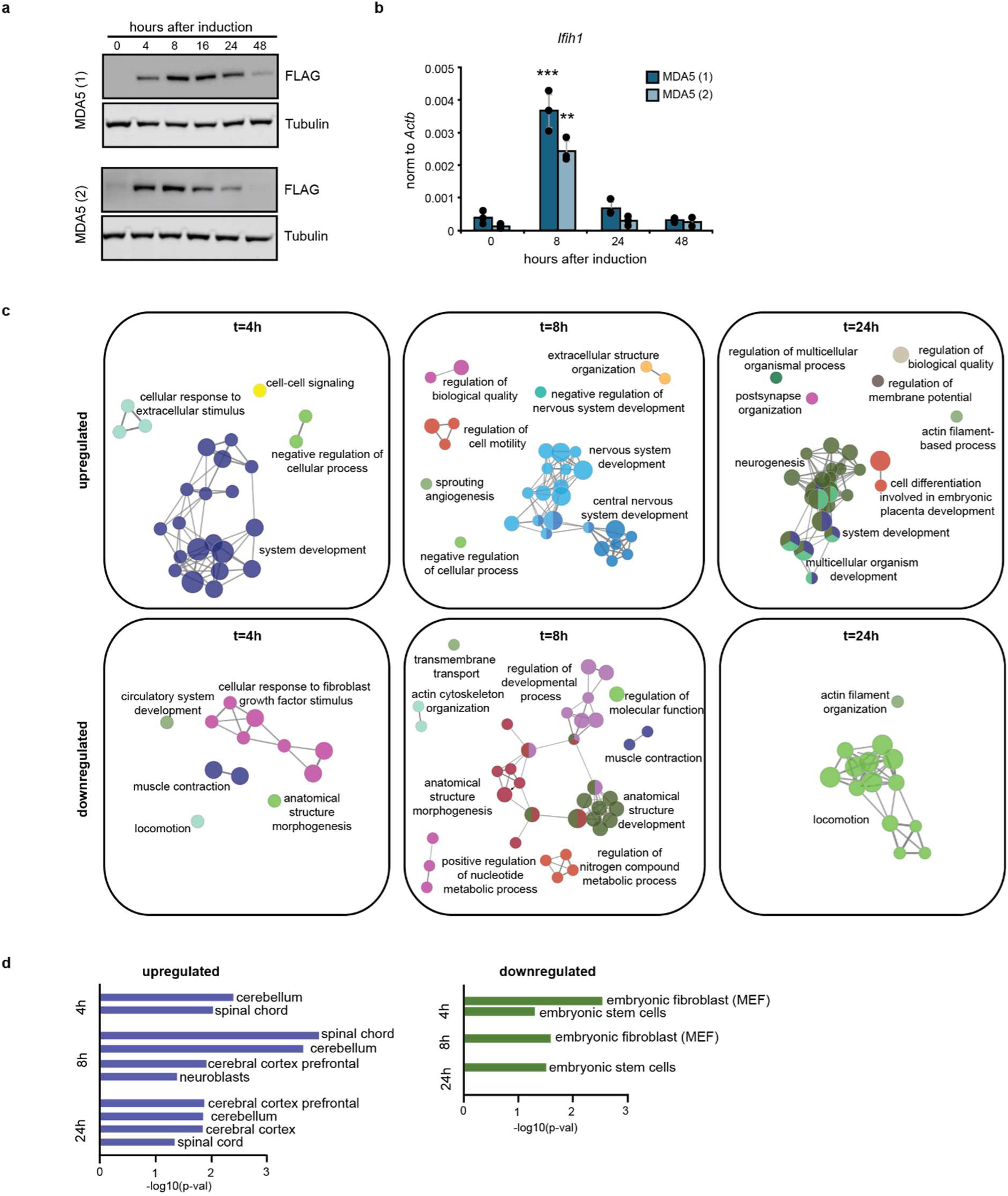
*Ifih1* results in defects of the pluripotency and differentiation programme of ESCs. (**a**) Two different ESC clones expressing a doxycycline-inducible N-terminal FLAG-tagged *Ifih1* gene were selected for further studies. Upon doxycycline stimulation, *Ifih1* (MDA5) expression was confirmed by western blot with anti-FLAG antibody. Tubulin serves as a loading control and expression was confirmed by RT-qPCR of *Ifih1* (**b**). Data represent the average of three biological replicates ± SD. Single factor ANOVA was used to calculate significant differences amongst comparisons, followed by an F-test for variance and appropriate two-tailed t-test, (*) p-val≤0.05, (**) p-val≤0.01, (***) p-val≤0.001. (**c**) Differentially upregulated and downregulated genes (abs log2FC>0.4, p-val≤0.05), in time-course analyses of *Ifih1* induction by total RNA high-throughput sequencing shared in both MDA5 expressing ESC clones (as shown in panel (**a**)) were used for gene ontology analyses (biological process) (**d**) The same set of shared up- and downregulated genes for both clones were analysed using the Mouse Gene Atlas for predicted cell types.

### MDA5 expression results in the loss of pluripotency factors and activation of immune pathways in ESCs

Considering the alterations in gene expression observed upon MDA5 induction, we hypothesise that these could be driven by epigenetic changes, including chromatin accessibility and/or differential activity of transcription factors. To test this possibility, we first assessed changes in the chromatin architecture upon MDA5 induction. We performed an Assay for Transposase-Accessible Chromatin coupled to high-throughput sequencing (ATAC-seq) and confirmed that MDA5 induces changes in chromatin accessibility, with 171 sites being more accessible, and a total of 382 losing accessibility after only 24 hours of MDA5 induction (Fig. 3a and Extended data Fig. 3, Supplementary Excel file 3). Gene Ontology analysis revealed that regions with more accessibility were associated with the nervous system function, similarly to the RNA high-throughput sequencing analyses, but also deleterious pathways such as apoptosis. On the other hand, regions losing accessibility were associated with signalling pathways regulating the pluripotency of stem cells, including TGF-β, Apelin and p53 (Fig. 3b). Interestingly, regions changing accessibility were enriched for transcription factor binding sites of pluripotency factors, including Oct4, Nanog, and several Sox and Klf family members (Extended data Fig. 3b). These findings were confirmed at the transcriptional level using the RNA-seq dataset. Using the loss-of-function (LOF) tool from the ShinyGO package (^44^), we found that the gene expression changes upon MDA5 induction were predicted to be driven, again, by transcription factors involved in pluripotency maintenance, such as Nanog, Oct4 (Pou5f1), and members of the Polycomb repressive complexes 1 and 2 (PRC1 and PRC2), including Suz12, Eed and Rnf2 (Fig. 3c, for full list see Extended data Fig. 3d). The PRCs play key roles in early development, by repressing the expression of lineage-specific markers during pluripotency and silencing the expression of pluripotency markers upon differentiation (^66,67^). In agreement with these observations, we confirmed that expression of the pluripotency factors *Nanog*, *Klf4* and *Pou5f1* was downregulated upon MDA5 induction. In contrast, *Sox2* expression increased (Fig. 3d). *Sox2* has been shown to increase during neuroectodermal differentiation, which agrees with the neuronal gene expression phenotype that ESCs acquire after MDA5 induction (Fig. 2d) (^68,69^). Altogether, our combined analyses suggest that the function and expression of transcription factors essential for early development are affected by MDA5.

**Fig. 3.**
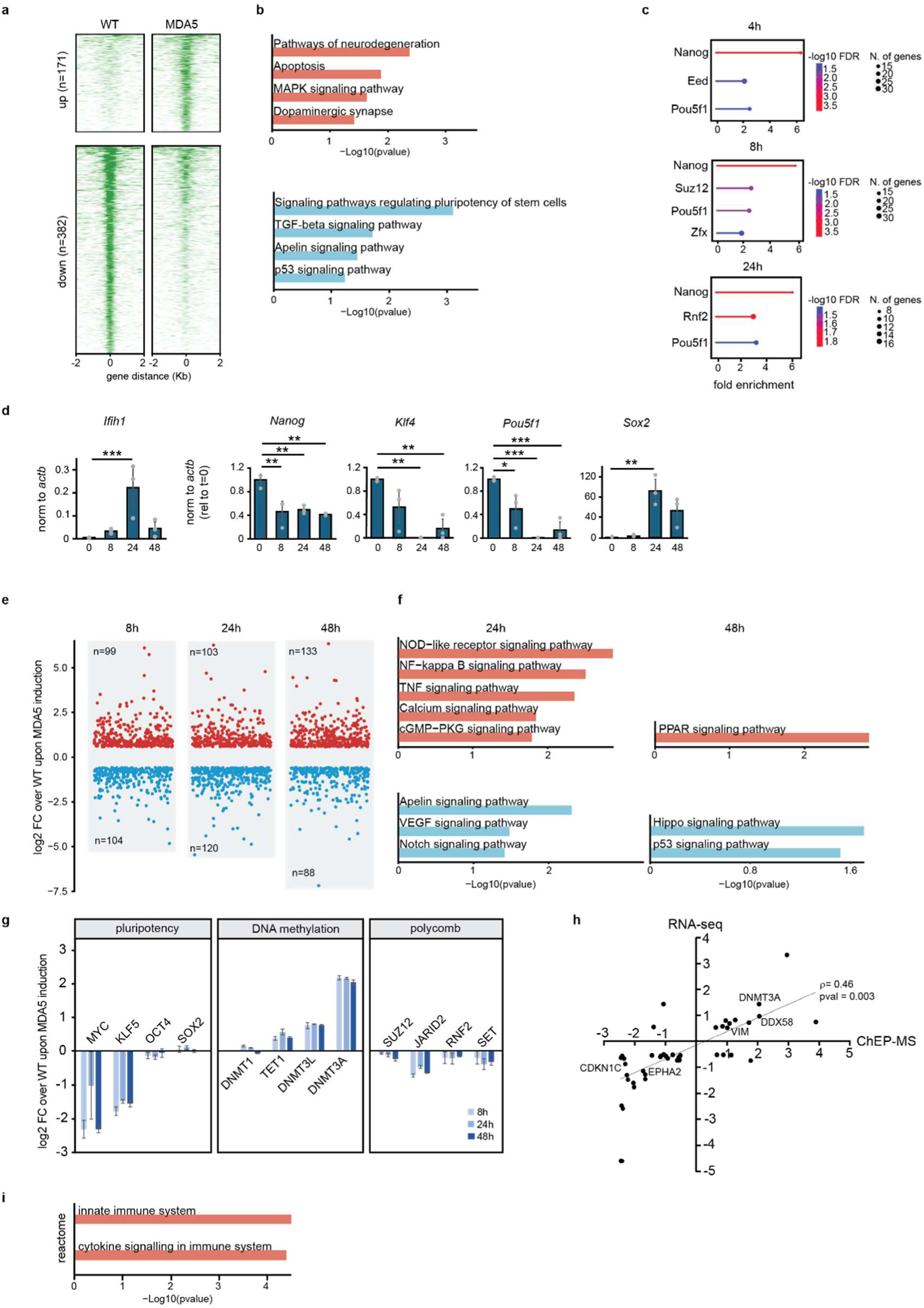
MDA5 expression leads to pluripotency transcription factor downregulation and innate immune signalling. (**a**) ATAC-seq data for WT and MDA5 expressing cells, both treated with doxycycline for 24 hours, showing changes in accessibility upon MDA5 expression. Data represent the average of two biological replicates. (**b**) KEGG pathway analyses of genes associated with differential ATAC peaks, with the more accessible (up, top) and less accessible (down, bottom) regions upon MDA5 expression. (**c**) Differentially expressed genes (RNA-seq) at t=4, 8 and 24h after *Ifih1* induction were used to compute significant similarities with loss-of-function (LOF) datasets for transcription factors (TFs) in mESCs. Only LOF-TFs from mESCs are included (for complete list see Extended data Fig. 3d). (**d**) RT-qPCR analyses of pluripotency genes during *Ifih1* induction. Data are the average of three biological replicates ± SEM. Single factor ANOVA was used to calculate significant differences amongst comparisons, followed by an F-test for variance and appropriate two-tailed t-test, (*) p-val≤0.05, (**) p-val≤0.01, (***) p-val≤0.001. (**e**) ChEP-MS analyses of a time course of *Ifih1* induction with the number of significantly different proteins over WT cells indicated (two-sided Student’s *t*-test, *P* < 0.05, FC > 2, 2 replicates). (**f**) KEGG pathway analyses for significantly differentially enriched chromatin-associated proteins obtained from ChEP-MS (two-sided Student’s *t*-test, *P* < 0.05, FC > 2) at 24h and 48h. (**g**) Comparison of the chromatin association of key regulators of pluripotency, DNA methylation and polycomb repressive complex between WT and MDA5 overexpressing mESCs (*n* = 2. Data are presented as the average ± s.e.m.) (**h**) Positive correlation between protein levels (ChEP-MS 0-48h) and RNA steady-state levels (0 to 8h). Spearman rank correlation coefficient; ρ=0.46, *p=0.003*. (**i**) Reactome analysis of correlating genes and proteins as identified in (**h**).

To provide further evidence of transcription factor dysregulation at the protein level and expand on the mechanism, we performed Chromatin-Enrichment for Proteomics followed by Mass Spectrometry (ChEP-MS). This approach provides a global overview of the chromatin composition, and how these changes under different conditions (^45,70^). The chromatin-associated proteome underwent substantial changes after 8 hours of MDA5 induction, with comparable numbers of proteins showing increased or decreased presence (absLog2FC>1, p < 0.05). Similar patterns were also observed at 24 and 48 hours after MDA5 induction (Fig. 3e, Supplementary Excel file 4). KEGG pathway analyses revealed that enriched proteins in chromatin were associated with immune and pro-inflammatory signalling pathways after MDA5 induction at 24 hours (Fig. 3f), while at 48 hours, pathways involved in shutting down pro-inflammatory and IFN responses were activated (^71,72^). On the other hand, factors involved in Notch, Hippo and p53 signalling, which are important for normal development, were depleted from the chromatin fraction after MDA5 induction (Fig. 3f). These results were reminiscent of the pathways obtained by ATAC-seq (Fig. 3b) and KEGG analysis of the time course RNA-seq data (Extended data Fig. 2b). Finally, we exploited these datasets to better understand the dynamics of key chromatin-associated proteins for pluripotency maintenance, including transcription factors and epigenetic regulators. Oct4 and members of the Myc and Klf families, as well as PRC components, were less enriched in chromatin fraction upon MDA5 induction. In contrast, factors involved in *de novo* DNA methylation were chromatin-enriched upon MDA5 induction (Fig.3g). To further probe for the correlation between the RNA-seq timecourse and ChEP-MS analyses, an early RNA-seq timepoint and a later ChEP-MS timepoint were selected, as we expect changes in RNA steady-state levels to result in changes at the protein level at a later timepoint. Using the genes and proteins shared in both datasets we found a significant positive correlation (p = 0.003, ρ=0.46) between RNA and protein levels, suggesting that the observed changes are not solely due to alterations in the subcellular localization of chromatin factors, but also reflect shifts in the gene expression profile of these cells (Fig. 3h). Interestingly, this same group of shared genes are associated with innate immune signalling, suggesting that expression of MDA5 in ESCs results in innate immune activation, and a concomitant loss of pluripotency accompanied by epigenetic and transcriptional reprogramming (Fig. 3i).

### Expression of MDA5 results in type I IFN activation upon recognition of endogenous dsRNAs

Both the high-throughput proteomic and RNA analyses suggested that MDA5 expression results in immune activation in ESCs in the absence of viral infection. To confirm this finding, we tested the expression levels of the type I IFN *Ifnb1*, *Tnfa*, and the ISGs *Isg15* and *Ifitm1* in MDA5-induced cells using RT-qPCR. MDA5 induction resulted in the expression of type I IFNs and ISGs in the absence of viral infections (Fig. 4a). ISG induction was also confirmed on the RNAseq data from the two independent MDA5 overexpressing clones (Extended data Fig. 4a). To test the importance of dsRNA recognition in activating the IFN response during MDA5 expression, we generated ESCs that express an MDA5 mutant lacking the ability to recognise dsRNA. The C-terminal domain (CTD, residues 898-1020) of MDA5 is proposed to play a key role in the initial binding to, and oligomerization on dsRNA (^73,74^) and based on this, an MDA5 variant lacking the CTD was generated (ΔCTD). Using *in vitro* pull-down assays, we confirmed that full-length MDA5 binds dsRNA (poly(I:C)), but not ssRNA (poly(C)), while the ΔCTD mutant failed to bind both dsRNA and ssRNA (Fig. 4b). We generated ESC lines stably expressing the ΔCTD mutant and found that expression of this truncated MDA5 form was not repressed as observed for full-length WT MDA5. In addition, this mutant did not activate the type I IFN response and consequent ISG expression, confirming that binding to endogenous dsRNA is required for ESCs to activate the IFN response upon MDA5 expression (Fig. 4c). Based on these results, we tested whether ESCs accumulate endogenous dsRNA by performing flow cytometry with antibodies against dsRNA (J2). As a control for staining specificity, we pre-treated cells with RNAseIII, a dsRNA specific endonuclease, before probing with J2 antibodies. We confirmed that dsRNA accumulated in ESCs and this signal was RNAseIII sensitive (Fig. 4d,e, Extended data Fig. 4b). On the other hand, the somatic IFN-competent microglial cell line, BV2, did not accumulate any RNAseIII-sensitive dsRNA (Fig. 4d,e). To further confirm the presence of dsRNA in ESCs we performed an immunoprecipitation (IP) using the dsRNA-specific antibody J2 followed by RNA high-throughput sequencing. We first looked at changes in biotypes of the enriched dsRNA compared to the non-enriched genes, and found an increase in the proportion of reads originating from protein-coding genes, a shift that has been observed before in dsRNA sequencing experiments (^75^) (Fig. 4f). Next, we calculated the minimum free energy (mfe) of the same set of enriched genes and found a significant decrease compared to all non-enriched genes, confirming the presence of more structured conformations, possibly dsRNA (Fig. 4g). Both sets of genes have similar GC content, indicating that the lower mfe is not merely the consequence of a higher GC content (median %GC 49.1 in non-enriched control, 48,9% in enriched dsRNAs) (Fig 4h). Interestingly, the enriched genes are longer than the controls (2866 vs 1894 median nts) (Fig. 4i). The main contributors of the lower mfe are the 3’ UTR and CDS (Fig4 j-l). In agreement with the proposed roles for TEs in forming dsRNA in mammalian cells (^14^), we found that the enriched dsRNAs harbour significantly more TEs per gene, suggesting that TE-sequences also contribute to dsRNA formation in early mouse development (Fig. 4m). These findings indicate that ESCs provide a permissive environment for dsRNA accumulation. However, this permissiveness is broken upon introduction of the dsRNA sensor MDA5, which activates the type I IFN response after dsRNA recognition.

**Fig. 4.**
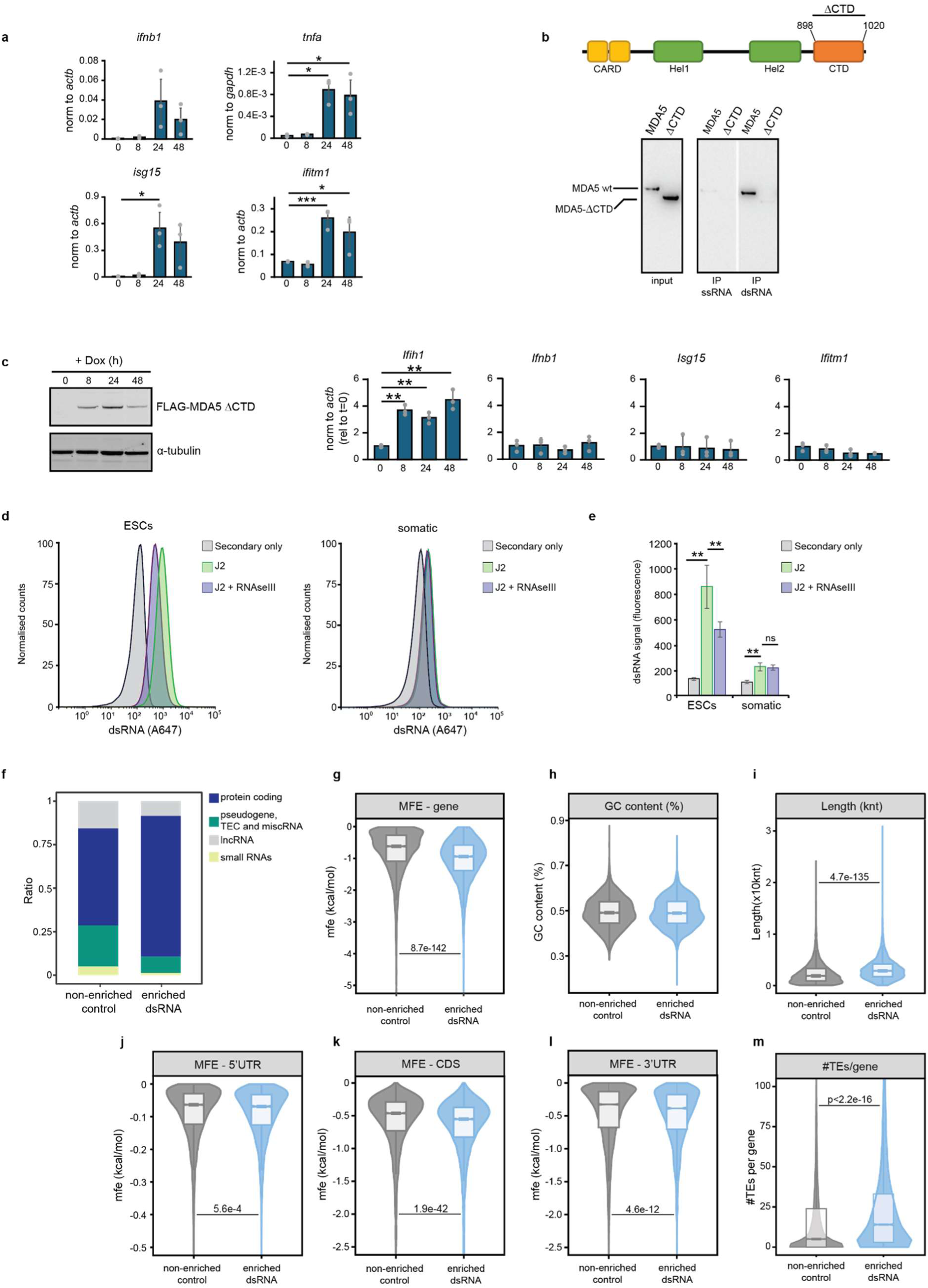
MDA5 expression results in IFN activation through endogenous dsRNA recognition. RT-qPCR analyses of the type-I IFN *Ifnb1*, pro-inflammatory gene *Tnfa* and ISGs, *Isg15* and *Iftim1* upon MDA5 induction time course. For *Ifih1* induction levels, see Fig. 3d. Data are the average of three biological replicates ± s.e.m., Single factor ANOVA was used to calculate significant differences amongst comparisons, followed by an F-test for variance and appropriate two-tailed t-test (*) p-val≤0.05, (**) p-val≤0.01, (***) p-val≤0.001. (**b**) Schematic representation of MDA5 domains, CARD (caspase activation and recruitment domain), Hel1 (Helicase ATP-binding), Hel2 (Helicase C-terminal), C-terminal domain (CTD from residue 898 to 1020) (top). Western blot analyses of pull-downs performed with full-length (WT) and truncated (ΔCTD) MDA5 using ssRNA (poly(C)) and dsRNA (poly(I:C)) (**c**) Expression of MDA5-ΔCTD in ESCs was confirmed by western blot analyses using anti-FLAG antibodies where tubulin serves as a loading control (left panel). RT-qPCR analyses of the same ESCs expressing doxycycline-inducible MDA5-ΔCTD cells confirmed expression of MDA5-ΔCTD and showed no *Ifnb1*, *Isg15* or *Ifitm1* expression upon induction. Data are the average of three biological replicates ± SEM, Single factor ANOVA was used to calculate significant differences amongst comparisons, followed by an F-test for variance and appropriate two-tailed t-test (*) p-val≤0.05, (**) p-val≤0.01, (***) p-val≤0.001. (**d**) Flow cytometry analyses of endogenous dsRNA accumulation using dsRNA specific antibodies (J2), in ESCs (left panel), and differentiated somatic BV2 cells (right panel). J2 + RNAse III, and secondary only samples serve as controls for antibody specificity and background, respectively. (**e**) Quantification of average dsRNA signal (n=3, ± SD). Single-factor ANOVA was used to calculate significant differences amongst comparisons, followed by an F-test for variance and appropriate two-tailed t-test. (**f**) Analysis of dsRNA immunoprecipitations by high-throughput sequencing shows a shift towards more protein-coding transcripts compared to the non-enriched fraction of the whole transcriptome. (**g**) Genes that were enriched in the dsRNA IP showed a significant reduction in minimum free energy, suggesting the presence of RNA with a more structured confirmation. (**h**) These same enriched genes had similar GC content, but were generally longer (**i**). (**j,k,l**) The major contribution to the lower mfe were found in the 3’UTR and CDS regions, rather than 5’UTRs. (**m**) Enriched genes had significantly more predicted TEs in their transcripts compared to the non-enriched fraction of the whole transcriptome.

### Who interferes with pluripotency: IFN production or IFN signalling?

Our results lead us to hypothesise that IFN production is responsible for the dysregulation of pluripotency genes. To test this possibility, genes downstream of MDA5 signalling were depleted to test their effect in perturbing pluripotency markers expression. Only when IRF3 and MAVS were depleted, and no IFN response was activated, pluripotency genes remained unchanged after MDA5 induction. On the other hand, depletion of IRF7 mimicked control cells, showing decreased expression of pluripotency genes with MDA5 (Fig. 5a, Extended data Fig. 5a). This suggest that ultimately the activation of IRF3 and possibly the production of IFN is responsible for the loss in the expression of pluripotency markers.

**Fig. 5.**
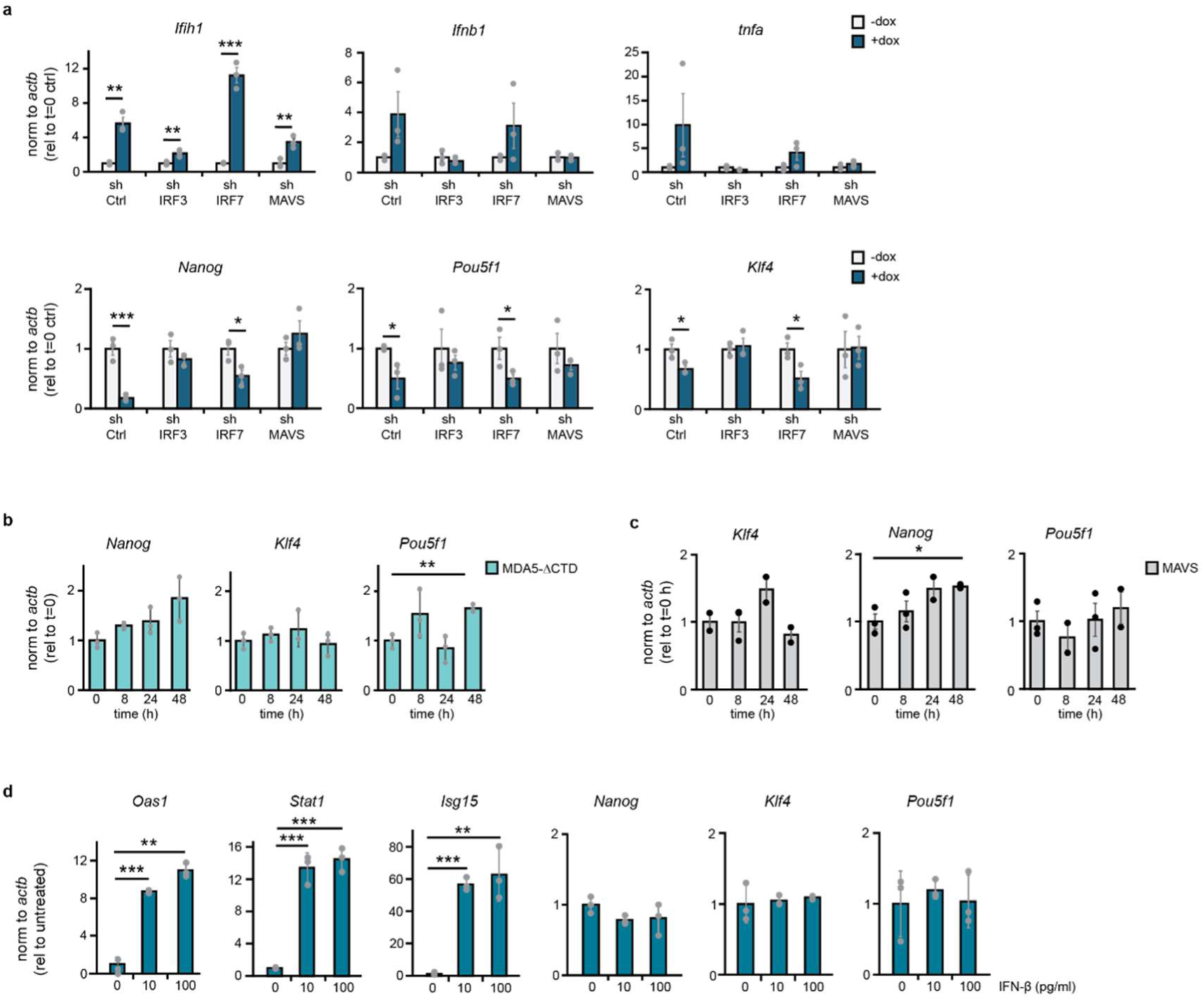
IRF3 activation and IFN production is responsible for pluripotency perturbation. (**a**) RT-qPCR analyses for the MDA5 gene *ifih1*, type-I IFN *Ifnb1*, pro-inflammatory gene *Tnfa* and pluripotency markers *Nanog*, *Pou5f1* and *Klf4* on MDA5 overexpressing cells depleted from *Irf3*, *Irf7*, *Mavs*, or shCtrl (ctrl). Data are the average of three biological replicates ± SEM, Single factor ANOVA was used to calculate significant differences amongst comparisons, followed by an F-test for variance and appropriate two-tailed t-test (*) p-val≤0.05, (**) p-val≤0.01, (***) p-val≤0.001. Depletion levels of *Irf3*, *Irf7* and *Mavs* were measured using RT-qPCR (Extended data Fig. 5a). (**b**) RT-qPCR analyses of pluripotency factors *Nanog*, *Pou5f1* and *Klf4* upon MDA5-ΔCTD expression. MDA5 levels are shown in Fig. 4c. Data are the average of three biological replicates ± SEM, Single factor ANOVA was used to calculate significant differences amongst comparisons, followed by an F-test for variance and appropriate two-tailed t-test (*) p-val≤0.05, (**) p-val≤0.01, (***) p-val≤0.001. (**c**) RT-qPCR analyses of pluripotency and IFN genes upon MAVS overexpression. Data are the average of three biological replicates ± SEM, Single factor ANOVA was used to calculate significant differences amongst comparisons, followed by an F-test for variance and appropriate two-tailed t-test (*) p-val≤0.05, (**) p-val≤0.01, (***) p-val≤0.001. (**d**) RT-qPCR analyses of ISGs and pluripotency genes upon exogenous IFN-β stimulation using two different concentrations (10 and 100 pg/ml). Data are the average of three biological replicates ± SEM, Single factor ANOVA was used to calculate significant differences amongst comparisons, followed by an F-test for variance and appropriate two-tailed t-test (*) p-val≤0.05, (**) p-val≤0.01, (***) p-val≤0.001.

To further confirm that pluripotency dysregulation is specific to MDA5 and IFN production, we measured the expression of pluripotency genes in MDA5-ΔCTD cells and MAVS overexpressing cells, which do not activate IFN production on its own (Fig. 4c and Extended data Fig. 5b). Neither of these resulted in perturbation of pluripotency genes (Fig. 5b,c). It therefore seems that the production of IFN is responsible for perturbing the pluripotency-associated gene expression programme. Finally, to test whether exogenous type I IFN stimulation could lead to a similar perturbation, ESCs were incubated with different amounts of IFN-β. This resulted in increased expression of ISGs such as *Oas1*, *Stat1* and *Isg15* by RT-qPCR, however, no significant changes in the expression of the pluripotency factors *Nanog*, *Klf4* or *Pou5f1* was observed (Fig. 5d). It therefore seems that IRF3-mediated production of IFN, and not the sensing of exogenous IFN, is responsible for perturbing the pluripotency-associated gene expression programme.

### Suppression of the dsRNA sensing pathway is conserved in other vertebrates

Both mouse and human embryonic stem cells fail to produce IFNs when challenged with dsRNA, suggesting that the suppression of this pathway is conserved across mammals (^30^). Jawed vertebrates, including fish and birds, also use the type I IFN response as the primary mechanism to defend against viruses. In zebrafish, the adaptive immune system is not well developed until 4 weeks post-fertilisation, making it a useful model to study the innate antiviral response in the early stages of development (^76–78^). Besides this, zebrafish are a powerful *in vivo* tool to assess the long-term effects of an active IFN response in early development on body plan formation. First, we confirmed that the expression of most RLR-signalling genes, including RIG-I, MDA5 and MAVS were also developmentally regulated in zebrafish (Fig. 6a) (^39^). Next, we investigated whether this developmentally regulated expression correlated with the ability of zebrafish to respond to dsRNA. To this end, 1-cell embryos were injected with the dsRNA analogue poly(I:C), and IFN production was measured by *in situ* hybridisation with probes against the ISG *Trim25*. Only after 24h did we observe an increase in *Trim25* staining (Fig. 6b) which was confirmed by analysing the expression of both *Trim25* and *Isg20* by RT-qPCR (Fig. 6c). DsRNA stimulation resulted in deleterious phenotypes in 60% of the injected embryos. Some embryos exhibited impaired head development (class 1), while others showed other defects including issues with trunk development (class 2) (Fig. 6d,e). These results suggest that activating the IFN response in early zebrafish development can have negative consequences. We also tested the consequences of IFN production in the differentiation ability of mESCs, as a model of *in vitro* development. IFN activation resulted in failure of mESCs to efficiently silence the expression of the major pluripotency factors during the first hours of differentiation, and premature expression of differentiating markers, such as *Hand1* (Extended data Fig. 6a).

**Fig. 6.**
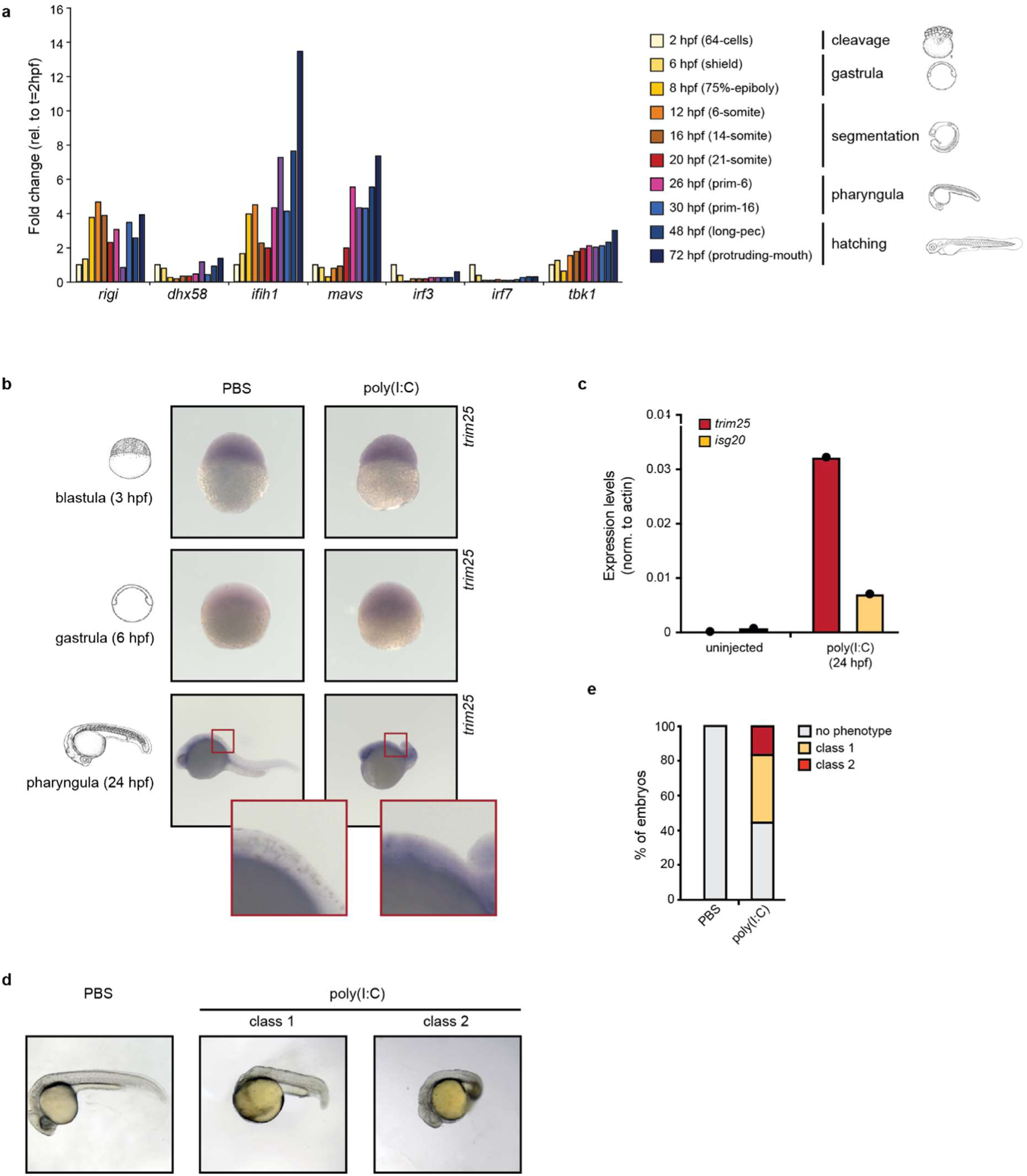
IFN is silenced in early zebrafish development and its activation leads to developmental defects. (**a**) Relative expression of RLR-signalling pathway genes across zebrafish development. Each timepoint is expressed relative to the expression at the earliest timepoint (2hpf). (**b**) *In situ* hybridization with probe against the ISG *trim25* to monitor IFN activation 3, 6 and 24h post-injection of dsRNA at the zygotic stage (0 hpf). PBS injection was used as a negative control (left). (**c**) RT-qPCR analyses of ISG expression at 24hpf after dsRNA stimulation with poly(I:C) using >50 embryos for each sample. (**d**) Phase-contrast microscopy of dsRNA-injected embryos (right), versus PBS-injected controls (left), 24 hours after injection. (**e**) Developmental defects were quantified as class 1 (failure to develop head), and class 2 (failure to develop the trunk) and represented as the proportion of embryos with defects vs non-defective.

All these together suggest that the ability to synthesise IFNs in zebrafish is developmentally regulated and, like mice, activation of the IFN response during early embryonic stages can have severe developmental consequences. We hypothesise that silencing of the dsRNA sensing pathway is conserved across jawed vertebrates.

## Discussion

The importance of silencing the IFN response during early development is illustrated by the observation that the RLR pathway, the primary activator of IFN production, is suppressed at different levels and through multiple mechanisms. Our findings show that in mESCs both the dsRNA sensor MDA5 and the transcription factor IRF7 are transcriptionally regulated during development, and increase their expression upon differentiation. Previously, overexpression of a catalytically active mutant form of IRF7 was also shown to result in IFN expression in ESCs, similar to MDA5 (^23^). It is still unclear what the mechanisms are that lead to silencing of MDA5 and IRF7 in ESCs. Our results show that neither DNA methylation nor Polycomb Repressive Complex 1 (PRC1) are required to silence MDA5 in ESCs (^79^). Interestingly, the mouse MDA5 gene harbours a CTCF binding site in the first intron, however, there is no evidence regarding the role of chromatin structure or loop extrusion on the expression of this gene during early development.

Transcriptional control is not the only mechanism silencing the IFN response in ESCs. We previously demonstrated that MAVS is post-transcriptionally silenced by miRNAs in mESCs. Removal of the MAVS-targeting miRNAs, or overexpression of MAVS restored the ability of mESCs to respond to dsRNA and increased their ability to defend from viruses (^22^). This redundancy in suppression of the IFN response by targeting different components of the pathway suggests that its downregulation is absolutely critical during early development. The suppression of the dsRNA-sensing pathway seems to be conserved across a range of species. Re-analyses of RNA high-throughput sequencing datasets demonstrate that other vertebrates also developmentally regulate the RLR pathway, including chickens (*Gallus gallus*), marmosets (*Callithrix jacchus*), macaques (*Macaca mulatta)*, frogs (*Xenopus tropicalis*) and zebrafish (*Danio rerio*) (^39,80–83^). In line with the dysregulation observed in mESCs, IFN induction in early development in zebrafish leads to developmental defects. Others have also observed that zebrafish embryos can produce IFN-φ1 upon dsRNA stimulation at 12 hours post-fertilization which is accompanied by a decrease in survival (^60^). These results together with our time course analysis suggest that the IFN response in zebrafish is enabled between 6 and 12 hours post fertilization, coinciding with increasing levels of MDA5 and once the maternal-to-zygotic transition and gastrulation have occurred. A more detailed time course analyses will be required to find when exactly zebrafish gain an active IFN response.

Our results suggest that it is important to silence MDA5 to prevent endogenous dsRNA recognition, and activation of the IFN response. The accumulation of dsRNA in ESCs was an unexpected finding, as normally, the presence of dsRNA is associated with viral infections. ESC-dsRNA derived from protein-coding genes with strong secondary structures and enriched in TE-sequences. Previous studies have implicated TE-derived sequences as substrates for viral nucleic acid sensors, including MDA5 (^14,84,85^). Interestingly, the overexpression of MDA5 in other cell types does not always result in spontaneous activation of the IFN response, and it seems to be a cell-line dependent phenotype, suggesting that cells accumulate different levels of unedited dsRNA (^86–91^). Future efforts will aim at understanding the function of these dsRNAs in ESC biology and development.

As a result of dsRNA recognition, MDA5 triggers the IFN response and a profound rewiring of the developmental programme of ESCs. The temporal scale of these changes suggests that it is driven by alterations in transcription factor activity, localisation and/or post-translational modifications and doesn’t rely on *de novo* protein synthesis. In agreement, ATAC-seq and ChEP-MS confirmed that MDA5 expression results in rapid changes in chromatin accessibility and chromatin-enrichment of transcription factors associated with pluripotency, differentiation, and immune responses. Interestingly, we also observed changes in the chromatin-enrichment of epigenetic factors, including DNA methyltransferases (DNMT) and polycomb-associated proteins. We hypothesise that some of the changes driven by MDA5 could be mediated by polycomb. For instance, MDA5 induction mimics some of the molecular defects observed in polycomb-deficient mESCs, where expression of differentiation markers occurs during pluripotency (^92^). MDA5-overexpressing ESCs also resemble PRC2-deficient mESCs (*Suz12^-/-^*), as they fail to efficiently repress pluripotency markers during differentiation (^93^). A more detailed study of which epigenetic and transcription factors are altered by MDA5 will provide further mechanistic insights into the role of this sensor in pluripotency and differentiation interference.

We also explored which specific factors and steps of the IFN response are responsible for altering the biology of ESCs. Our results suggest that it is not MDA5 itself who is interfering with pluripotency, but rather IRF3 activation and production or transcription of IFN itself. Similarly, overexpression of IRF7 also results in dysregulation of the ESC’s gene expression programme (^23^). In agreement, we only observed alterations in the pluripotency gene expression programme when we force cells to produce IFNs, but not upon sensing of exogenous IFN. This is possibly due to the necessity of embryos to still be able to respond to maternal IFNs during early development (^94^). Previous studies have also hinted towards interactions between pluripotency factors and the IFN response. For instance, the ability of IRF7 to induce the IFN response in ESCs can be reversed by overexpressing the pluripotency factors Oct4, Sox2 and Klf4 (^23^). It is unknown if this observation is only specific to IRF7, and the exact mechanism by which these TFs can block IRF7 activity. Interestingly, we observed downregulation of Nanog, Oct4 and Klf4 levels upon MDA5 induction, suggesting that changes in the expression of these factors could also be responsible for the dysregulation of the pluripotency gene expression programme.

Importantly, the production of IFN is not only toxic during early development. A strong IFN response during later stages can also lead to developmental abnormalities. Congenital infections, where maternal infections are vertically transmitted to the foetus, are a major cause of neurodevelopmental complications (^95^). Most infections occur post-implantation, affecting IFN-competent cells after evading maternal-placental defence systems. Similarly, interferonopathies—Mendelian diseases characterized by excessive IFN secretion—often lead to neurological developmental issues despite IFN activation occurring later in development (^13,96^). Given these patterns, it is reasonable to speculate that they share a common mechanistic basis with our findings in mESCs.

To conclude, we hypothesise that the gene expression program required for early development is incompatible with an active dsRNA-based antiviral response. Production of IFNs is deleterious for development and results in a global disruption of normal pluripotency and development, likely due to the incompatibility between the active transcription factors and available chromatin landscape. To avert this disruption, cells have suppressed the IFN response at multiple levels, with the unavoidable consequence of making them more susceptible to viral infections.

## Acknowledgements

This work was funded by the Wellcome Trust grants (221737/Z/20/Z and 107665/Z/15/Z), the Royal Society grant (RGS\R1\191368) and the Wellcome Trust iTPA (PIII021) to S.M. The Marks lab is supported by a Nederlandse Organisatie voor Wetenschappelijk Onderzoek (NWO) XL grant (OCENW.XL21.XL21.100) and a Radboud Science faculty grant (IRP voucher), ZL acknowledges support from the China Scholarship Council (CSC). C.R. is supported by EU funding under the MUR PNRR (Project no. E63C22001220001). T.T. is supported by AIRC under MFAG 2020 (ID. 24883 project) and by EU funding under the MUR PNRR ‘National Center for Gene Therapy and Drugs based on RNA Technology’ S6 RINGTAIL (Project no. CN00000041 CN3 RNA). We thank Robert Illingworth and Elias Friman for discussions on the epigenetic regulation of MDA5.

## Data availability

Bulk RNAseq of MDA5 induction, dsRNA IP, ATACseq and proteomics data are available under SRA Bioproject PRJNA1219136, PRJNA1223341, GEO database GSE289814 and PRIDE proteomics database PXD059977 respectively.

## Code availability

All custom code used for the analysis of data and generation of plots is available upon request

## Contributions

S.M. and J.W. conceived the project and wrote the manuscript with the help from J.L.W. A.I. performed computational analyses of RNA-sequencing datasets. H.M, Z.L. and L.M. designed, performed and analysed the ATACseq and ChEP-MS experiments. T.T. and C.R analysed the dsRNA data and A.A., P.G.M. and S.H. designed, performed and analysed the zebrafish experiments.

## Extended data figures

**Extended data Fig. 1.**
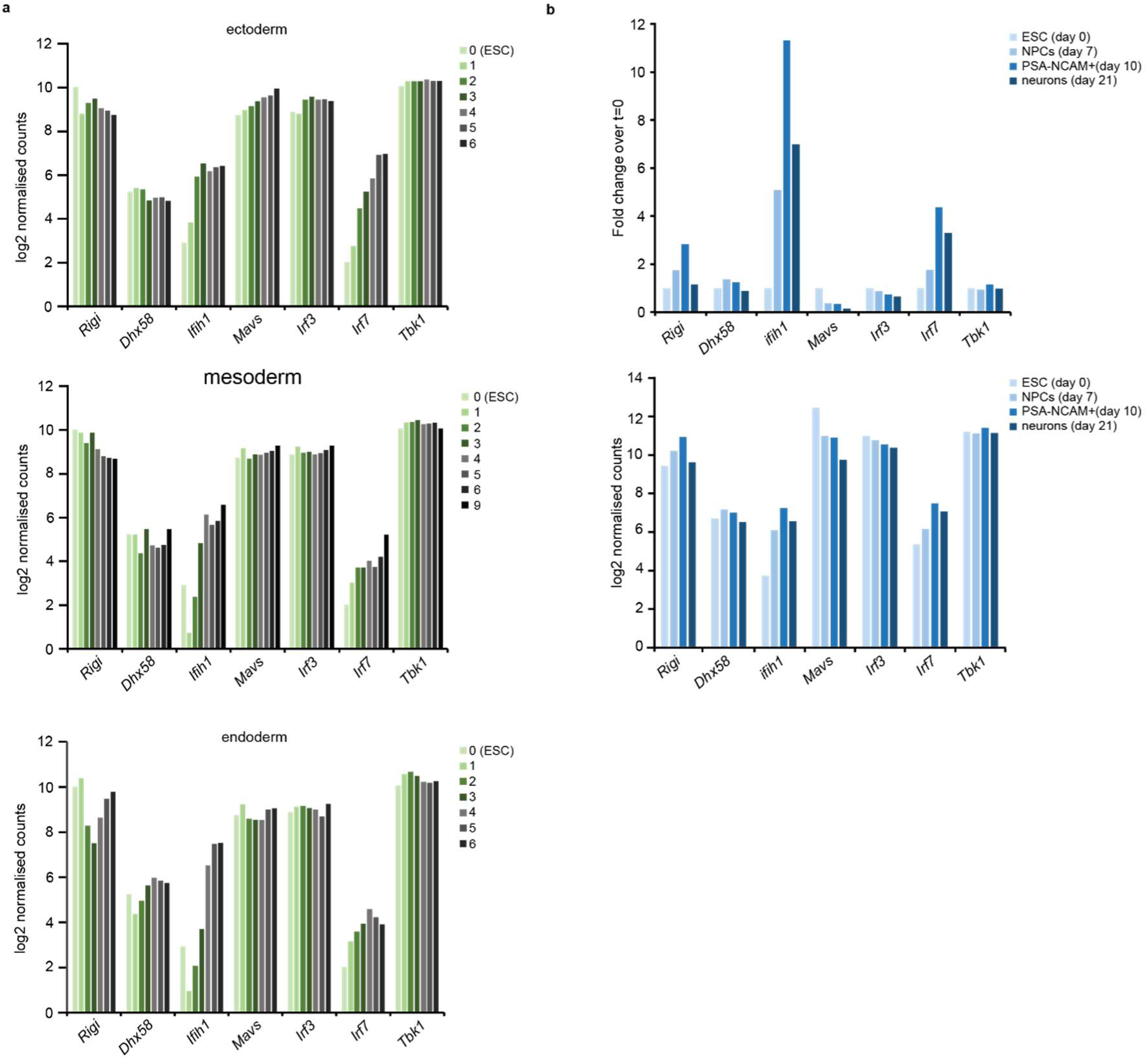
*Ifih1* expression is induced upon differentiation. **(a)** Absolute expression levels of genes involved in TLR signalling, RLR signalling pathway and JAK/STAT signalling analysed by high-throughput RNA sequencing of mouse embryonic stem cells (mESCs) differentiating to ectoderm (top), mesoderm (middle) and endoderm (bottom) (MTAB_4904, ^38^). Samples were taken at successive days after starting differentiation (day 0 corresponds to ESCs, until day 6 or 9 of differentiation). Expression is calculated as log2 normalised counts. **(b)** Relative (top) and absolute (bottom) expression levels of genes involved in RLR signalling in dataset of high-throughput RNA sequencing (GSE125413, ^37^). Cells were differentiated from ESC to neurons, with samples taken at days 0 (ESC), 7 (neural progenitor cells), 10 (PSA-NCAM+ cells) and day 21 (mature neurons). PSA-NCAM is a marker for intermediate differentiation. Expression is calculated as normalised counts relative to ESCs (day 0) for the upper panel and as log2 normalised counts for the lower panel.

**Extended data Fig. 2.**
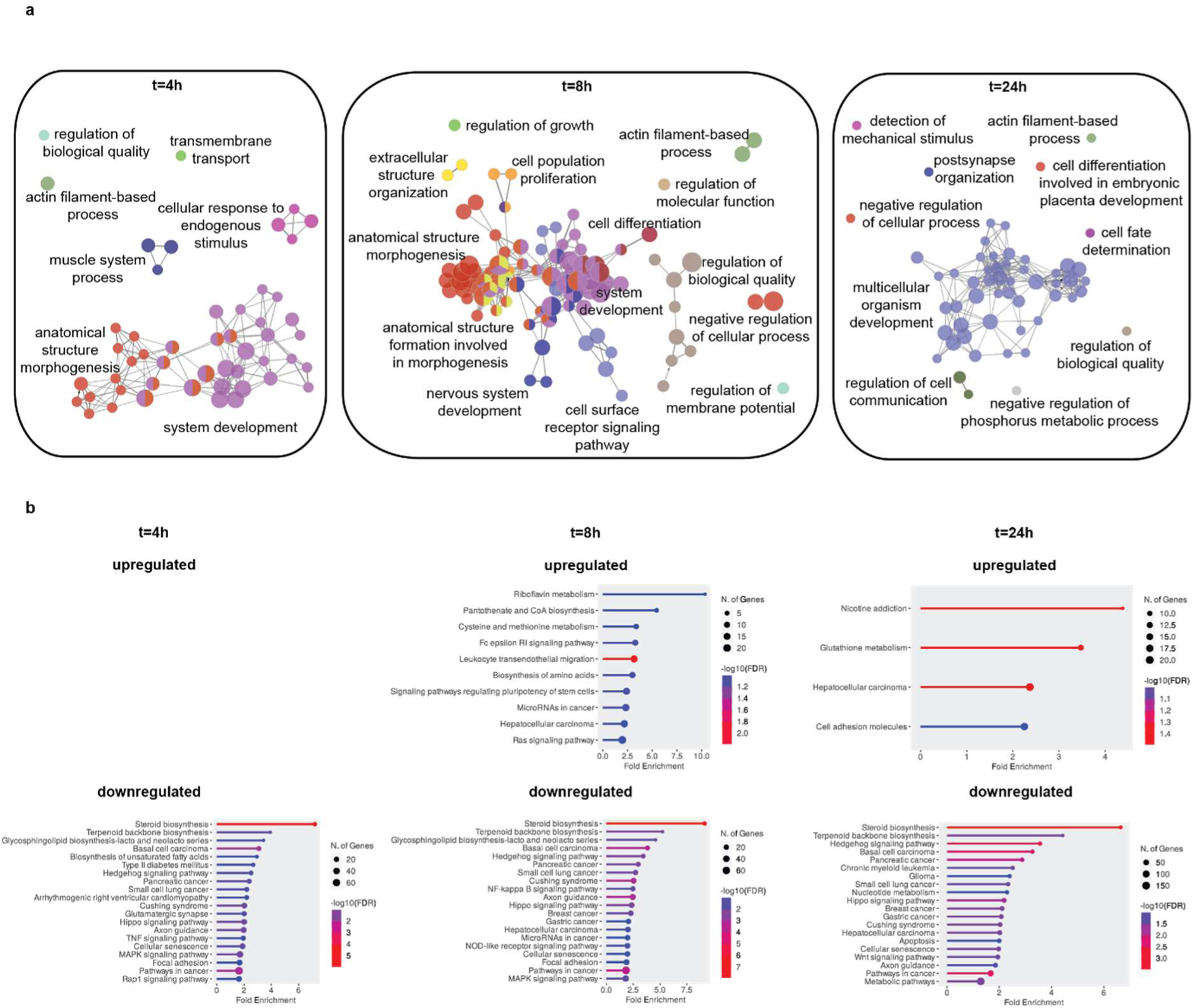
Time-course analyses during *Ifih1* induction by RNA-seq. (**a**) Differentially expressed genes (abs log2FC>0.4, p-val≤0.05) that were either up- or down regulated in both clones were used for gene ontology analyses (biological process). (**b**) KEGG analysis of same dataset as in (**a**).

**Extended data Fig. 3.**
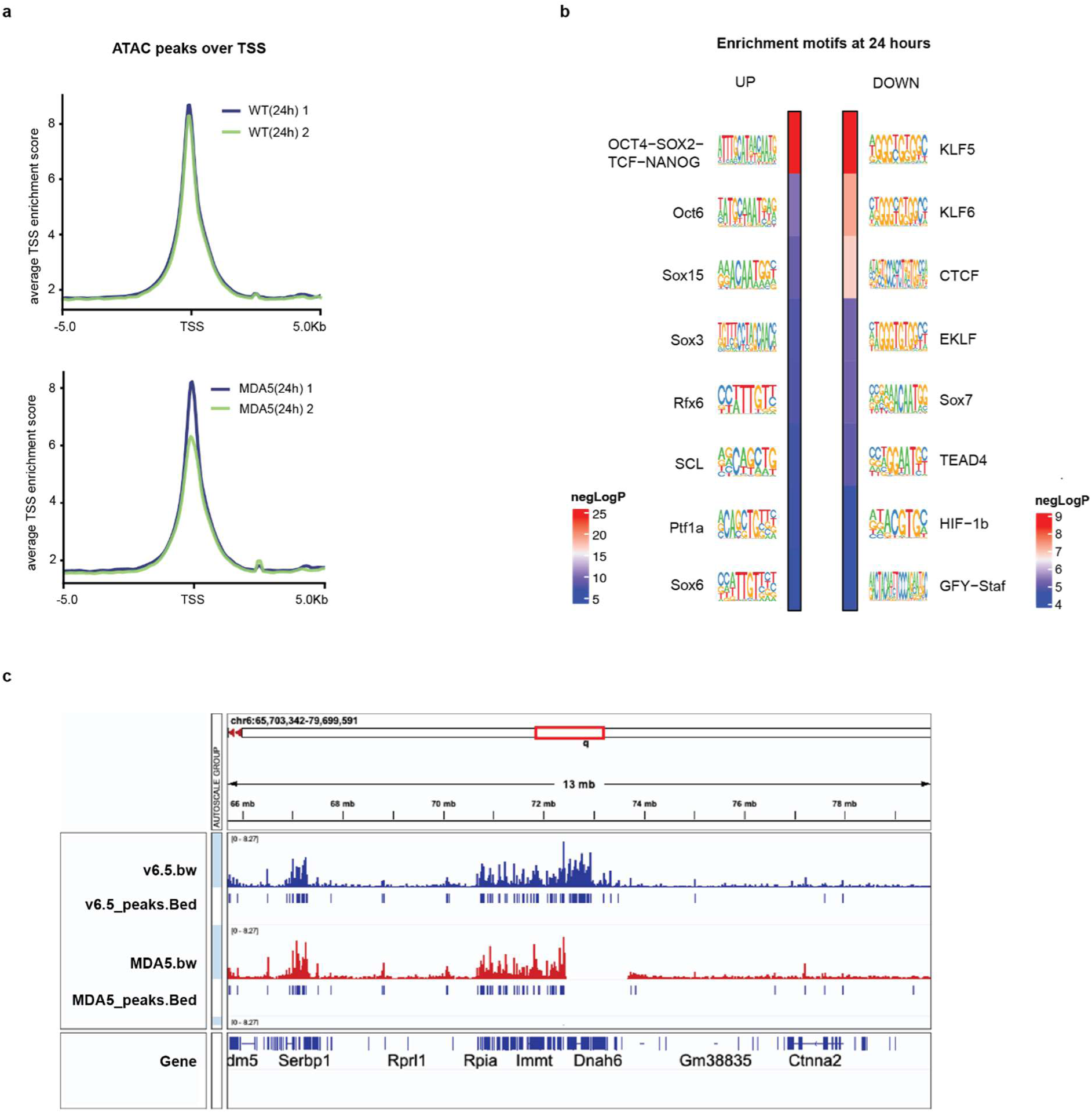
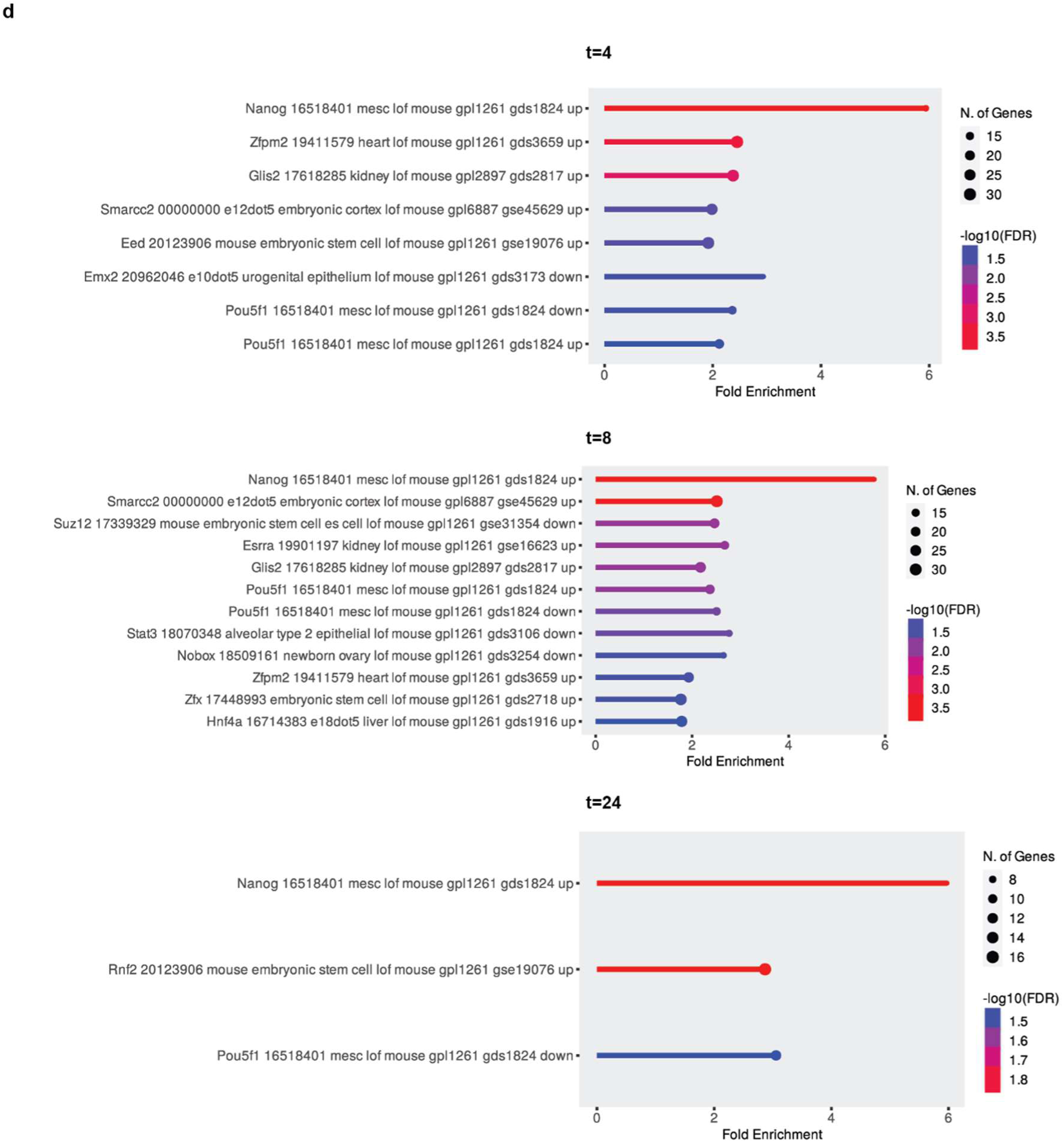
ATACseq peaks distribution and motif enrichment. (**a**) Feature distribution of ATAC peaks with mean read count frequency of peaks at the transcription start site (TSS) for the different experimental samples in WT (top) and MDA5-expressing ESCs (bottom). (**b**) Known motif analysis was performed on ATAC peaks that were up- or down-regulated at 24 hours compared to t=0 hours. Sequence weight matrixes of matched DNA-binding motifs are shown, with -log2(P value) represented by colour. (**c**) ATAC seq revealed a region missing MDA5 overexpressing clone 1, which was confirmed by RNAseq. The other clone (2) did not have the same deletion. (**d**) Differential gene expression at t=4, 8 and 24h after *Ifih1* induction was used to compute significant similarities with loss-of-function (LOF) datasets for transcription factors (TFs) in mouse.

**Extended data Fig. 4.**
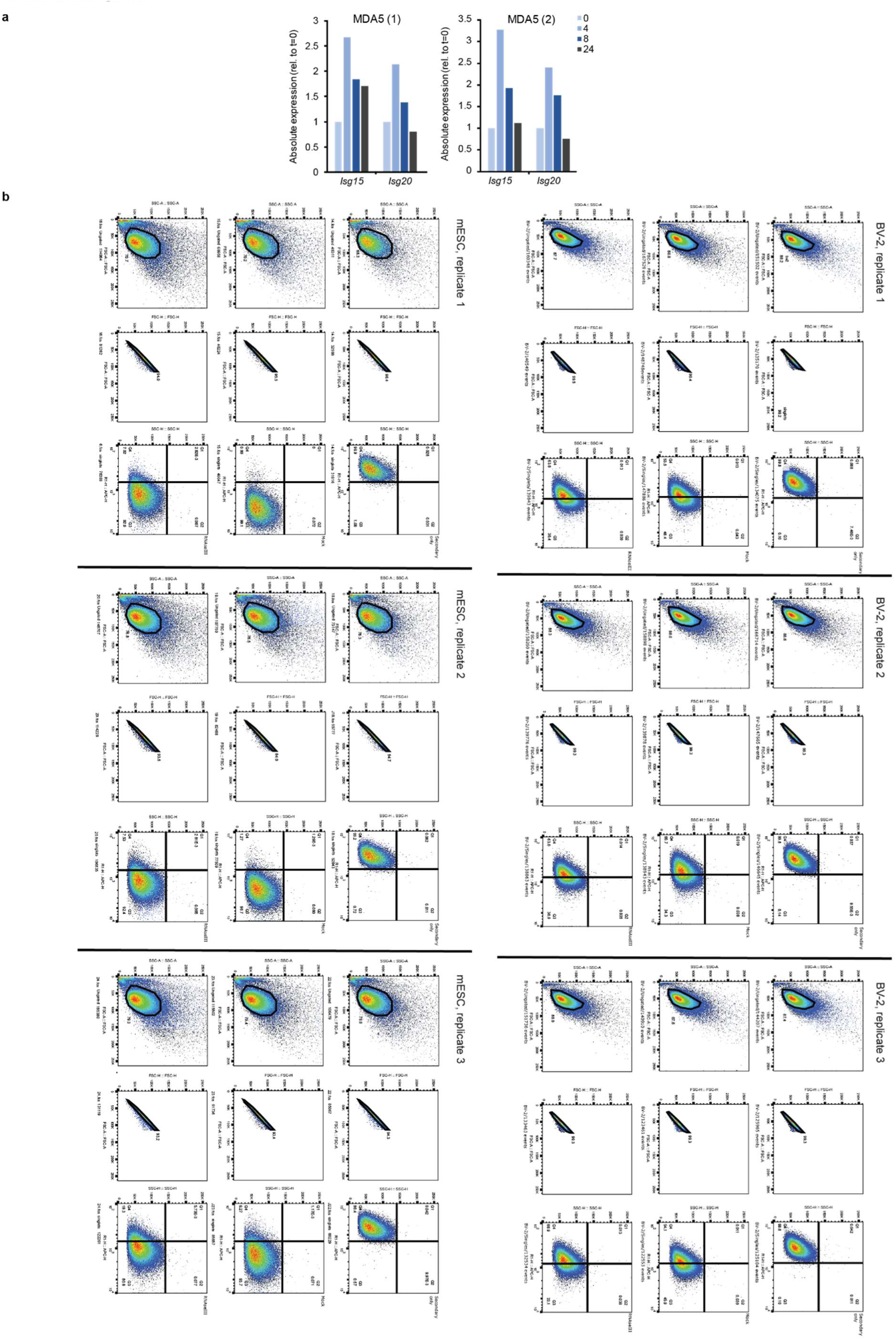
RNAseq data reveals induction of IFN response and Flow cytometry for dsRNA in mouse ESCs and BV2 cells. (**a**) Expression levels found in the RNAseq data of MDA5 overexpression in clones 1 and 2 revealed induction of ISGs and *Ifnb1* and *Tnf*. (**b**) Flow cytometry of three mouse ES- and BV2 cell replicates showing gating from main population of cells (left), gating for singlets (middle) and dsRNA signal using the appropriate channel (right).

**Extended data Fig. 5.**
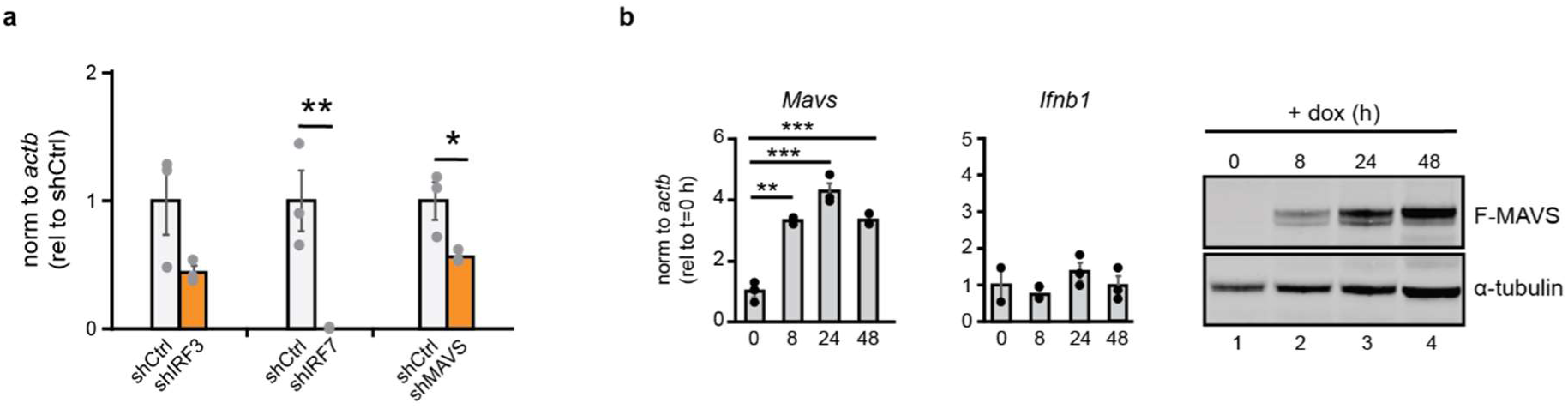
shRNA depletion levels. MAVS overexpression fails to induce IFN response. (**a**) RT-qPCR analyses for depletion levels of *Irf3*, *Irf7* and *Mavs*. Data are the average of three biological replicates ± SEM, Single factor ANOVA was used to calculate significant differences amongst comparisons, followed by an F-test for variance and appropriate two-tailed t-test (*) p-val≤0.05, (**) p-val≤0.01, (***) p-val≤0.001. (**b**) Time course of MAVS expression after induction in ESCs. RT-qPCR analysis showed sustained level of expression and no induction of the IFN response. Western blot analysis using anti-FLAG antibodies showed increasing protein levels. Tubulin was used as loading control.

**Extended data Fig. 6.**
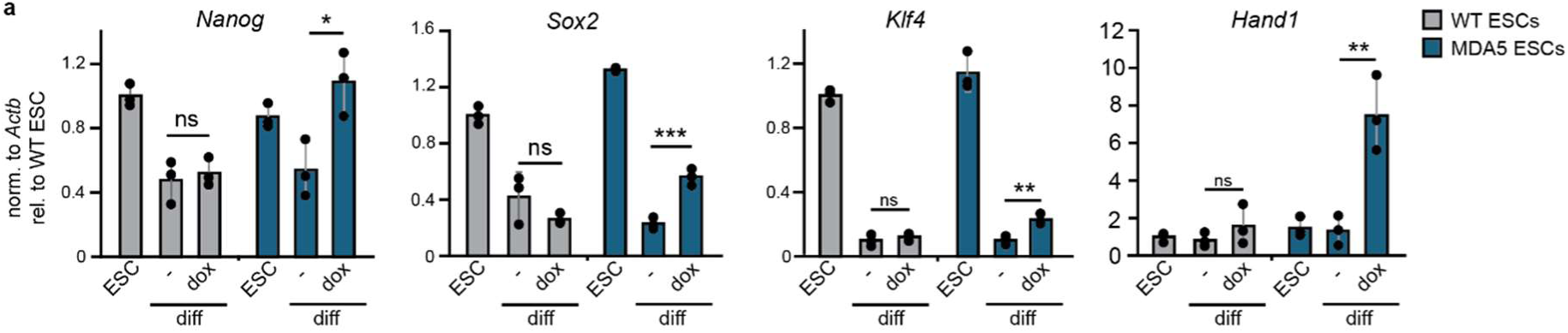
Differentiation of mESCs is dysregulated when MDA5 is overexpressed. **(a)** Both WT and MDA5 (clone 1) expressing ESCs were differentiated *in vitro* using embryoid bodies for 24 hours. Expression of pluripotency (*Nanog*, *Sox2*, *Klf4*) and differentiation markers (*Hand1*) were compared in untreated cells (ESCs), differentiating cells without doxycycline (-), and differentiating cells in the presence of doxycycline (dox). Data represent the average of three biological replicates ± SD. Single factor ANOVA was used to calculate significant differences amongst comparisons, followed by an F-test for variance and appropriate two-tailed t-test, (*) p-val≤0.05, (**) p-val≤0.01, (***) p-val≤0.001. Time-course analyses during *Ifih1* induction by total RNA high-throughput sequencing.

---

**Supplementary file 1**. Log2 normalised counts of MDA5 induction time course for MDA5 clone 1 (#10), MDA5 clone 2 (#2) and the parental line v6.5.

**Supplementary file 2**. List of the differentially expressed genes (abs log2FC>0.4, p-val≤0.05) that are shared between the two clones used, separated in up- and downregulated genes for each timepoint.

**Supplementary file 3.** Differentially detected peaks for ATAC seq of MDA5 clone 1 compared to parental line v65.

**Supplementary file 4.** ChEP-MS LFQ values for detected proteins in MDA clone 1 and parental line v6.5 (WT) for timepoint 0, 8, 24 and 48 hours post induction.

**Supplementary file 5**. Oligos used for RT-qPCR and cloning

